# The developmental changes in intrinsic and synaptic properties of prefrontal neurons enhance local network activity from the second to the third postnatal week in mice

**DOI:** 10.1101/2020.01.21.913913

**Authors:** Katerina Kalemaki, Angeliki Velli, Ourania Christodoulou, Myrto Denaxa, Domna Karagogeos, Kyriaki Sidiropoulou

## Abstract

The prefrontal cortex (PFC) is characterized by protracted maturation. The cellular mechanisms controlling the early development of prefrontal circuits are still largely unknown. Our study delineates the developmental cellular processes in the mouse medial PFC (mPFC) during the second and third postnatal weeks and characterizes their contribution to the changes in network activity. We show that spontaneous inhibitory postsynaptic currents (sIPSC) are increased while spontaneous excitatory postsynaptic currents (sEPSC) are reduced from the second to the third postnatal week. Drug application suggested that the increased sEPSC frequency in mPFC at P10 is due to depolarizing GABA_A_ receptor (GABA_A_R) function. To further validate this, perforated patch-clamp recordings were obtained and the expression levels of K-Cl co-transporter 2 (KCC2) protein were examined. The reversal potential of IPSCs in response to current stimulation was significantly more depolarized at P10 compared to P20 while KCC2 expression is decreased. Moreover, the number of parvalbumin-expressing GABAergic interneurons increase from P10 to P20 in the mPFC and their intrinsic electrophysiological properties significantly mature. Using computational modeling, we show that the developmental changes in synaptic and intrinsic properties of mPFC neurons contribute to the enhanced network activity in the juvenile compared to neonatal mPFC.

## Introduction

During early postnatal development, major events that contribute to cortical circuit maturation include spatial and temporal patterns of electrical activity, intrinsically determined cell death of early postnatal cortical interneurons and the depolarizing action of the neurotransmitter GABA (γ-aminobutyric acid) (Khazipov et al. 2004; Khazipov and Luhmann 2006; Allene et al. 2008; Brockmann et al. 2011; Ben-Ari et al. 2012; Southwell et al. 2012; Khazipov et al. 2013; Kirmse et al. 2015; Mòdol et al. 2019). The developmental switch of GABA action from depolarizing to hyperpolarizing results from changes in chloride co-transporter expression: Na^+^-K^+^-Cl^−^ cotransporter 1 (NKCC1), the Cl^−^ importer, is highly expressed early in development, while the expression of the co-transporter KCC2, the Cl^−^ exporter, increases after the first postnatal week (Ben-Ari 2001; 2002; Ben-Ari et al. 2007; Ben-Ari 2012). In addition, both intrinsic properties of neurons and synaptic transmission undergo dramatic changes during early postnatal development in a brain-area specific manner (Kriegstein et al. 1987; McCormick and Prince 1987; Burgard and Hablitz 1993; Ramoa and McCormick 1994; Bahrey and Moody 2003). Most studies on these developmental changes of the GABAergic system in the cortex have focused on the primary somatosensory cortex, visual cortex and hippocampus (Bartolini et al. 2013; Le Magueresse and Monyer 2013; Hensch 2016).

The barrel cortex (BC) is part of the primary somatosensory cortex and is organized vertically in columns of cells associated with sensory perception (Guo et al. 2014) and horizontally in six layers of distinct cell types. In contrast, cortical association areas such as the prefrontal cortex (PFC) regulate cognitive functions and do not directly control sensory information (Fuster 2015). Anatomically, the mouse medial PFC (mPFC) is defined as the agranular part of frontal lobe, lacking the prominent granular layer IV and is divided into distinct subregions, namely infralimbic, prelimbic and cingulate cortex (Heidbreder and Groenewegen 2003; Van De Werd et al. 2010). The timeline of mPFC development is delayed compared to other sensory cortices, such as BC (Casey et al. 2000; Best and Miller 2010; Kolb et al. 2012, Rinetti-Vargas et al., 2017). From infancy to adulthood, the developing mPFC undergoes considerable transcriptional, structural and functional changes (Diamond 2005; Tsujimoto 2008; Kolb et al. 2012; Schubert et al. 2014; Kroeze et al. 2017).

While adolescent development of prefrontal circuitry and the underlying cellular mechanisms have been addressed by a large number of studies, few investigations tackled the wiring processes at earlier stages (Brockmann et al. 2011; Bitzenhofer et al. 2015; 2017), in particular the mPFC between the second (neonatal) and third (juvenile) postnatal week. Here, we aim to fill this gap by investigating the synaptic and intrinsic properties of neonatal and juvenile mPFC neurons, with a primary focus on the GABAergic system, and identifying their contribution to the development of network activity.

## Methods

All *in vitro* experiments with mice took place under an experimental protocol approved by the Research Ethics Committee and by our Institutional Animal Care and Use Committee that has been approved by the Veterinarian Authorities Office (protocol license no. 93164). Experiments were carried out by trained scientists and in accordance with the 3R principles. *In vivo* experiments were performed in compliance with the German laws and the guidelines of the European Community for the use of animals in research and were approved by the local ethical committee (015/17, 015/18).

### Animals

The *in vitro* experiments were performed on male C57Bl/6J; Lhx6Tg(Cre); R26R-YFP+/+ mice from animal facility of IMBB-FORTH were used. For the *in vivo* experiments, timed-pregnant C57BL/6J mice from the animal facility of the University Medical Center Hamburg-Eppendorf were used. The day of vaginal plug detection was defined as embryonic day (E)0.5, whereas the day of birth was defined as P0. The offspring of both sexes are used for *in vivo* electrophysiology recordings. All procedures were performed according to the European Union ethical standards outlined in the Council Directive 2010/63EU of the European Parliament on the protection of animals used for scientific purposes.

Mice were housed with their mothers and provided with standard mouse chow and water ad libitum, under a 12 h light/dark cycle (light on at 7:00 am) with controlled temperature (21^◦^C). The P10 experimental group includes ages P9-P11 and the P20 group includes ages P19-P21, also referred to as second and third postnatal weeks or neonatal and juvenile, respectively. All efforts were made to minimize both the suffering and the number of animals used.

### *In vitro* extracellular recordings

#### Slice Preparation

Mice (P10 and P20) were decapitated under halothane anesthesia. The brain was removed promptly and placed in ice cold, oxygenated (95% O_2_ –5% CO_2_) artificial cerebrospinal fluid (aCSF) containing (in mM): 125 NaCl, 3.5 KCl, 26 NaHCO_3_, 1 MgCl_2_ and 10 glucose (pH = 7.4, 315 mOsm/l). The brain was blocked and glued onto the stage of a vibratome (Leica, VT1000S). Rostrocaudal coronal slices (400 μm thick) containing either the mPFC (prefrontal cortex) or the BC (barrel cortex) region were selected and transferred to a submerged chamber, which was continuously superfused with oxygenated (95% O_2_ –5% CO_2_) aCSF containing (in mM): 125 NaCl, 3.5 KCl, 26 NaHCO_3_, 2 CaCl_2_, 1 MgCl_2_ and 10 glucose (pH = 7.4, 315 mOsm/l) at room temperature (RT). The slices were allowed to equilibrate for at least 1 h in this chamber before recordings began. Slices were then transferred to a submerged recording chamber, continuously superfused with oxygenated (95% O_2_ –5% CO_2_) aCSF (same constitution as the one used for maintenance of brain slices) at RT during recordings.

#### Data Acquisition

Electrophysiological recordings were performed in all experimental groups under the same conditions as described below. Pulled glass micropipettes were filled with NaCl (2M) and placed in layers II/III of PFC and BC. Platinum/iridium metal microelectrodes (Harvard apparatus United Kingdom, 161 Cambridge, United Kingdom) were placed on layer II/III of the mPFC and the BC, about 300 μm away from the 1MΩ recording electrode, and were used to evoke field excitatory postsynaptic potentials (fEPSPs). Local field potentials (LFPs) were amplified using a headstage with selectable high pass filter of 30 Hz to remove any offsets coupled to a Dagan BVC-700A amplifier, amplified 100 times and low-pass filtered at 1-kHz. A notch filter was used to eliminate line frequency noise at 30 Hz. Signals were digitized using the ITC-18 board (InstruTech, Inc.) on a PC with custom-made procedures in IgorPro (Wavemetrics, Inc.) and stored on a PC hard drive. All voltage signals were collected at a sampling frequency of 100 kHz (fs = 100 kHz).

For evoked fEPSPs, the electrical stimulus consisted of a single square waveform of 100 μs duration given at intensities of 0.1– 0.3 mA (current was increased from 0.1 mA to 0.3 mA, with 0.1 mA steps) generated by a stimulator equipped with a stimulus isolation unit (World Precision Instruments, Inc.). The effect of GABAAR activation was investigated by bath application of 2 μM Diazepam (GABAAR agonist). Diazepam was acquired from the Pharmacy of the University General Hospital in Heraklion as a 5 mg/ml solution and was diluted in aCSF during recordings. Other drugs used include CNQX (10μM), AP5 (50μM) and bumetanide (10μM) (Tocris). For spontaneous activity recordings, 20 5-sec recordings were acquired without any stimulation.

#### Data Analysis

Data were analyzed using custom-written procedures in IgorPro software (Wavemetrics, Inc.). No additional high-pass filters were applied to the raw data. For evoked recordings, the peak values of the fEPSP were measured using the minimum value of the synaptic response (4–5 ms following stimulation) and were compared to the baseline value prior to stimulation. Both parameters were monitored in real- time in every experiment. A stimulus– response curve was then plotted using stimulation intensities between 0.1 and 0.8 mA, in 0.1 mA steps. For each different intensity level, two traces were acquired and averaged.

### *In vitro* patch-clamp recordings

#### Slice Preparation

Mice were decapitated under halothane anesthesia. The brain was removed immediately and coronal slices of mPFC and BC (300–350 μm thick), using a vibratome (Leica, VT1000S, Leica Biosystems) were prepared from mice at the ages of P10 and P20 in ice-cold oxygenated (95% O_2_ - 5% CO_2_) modified choline-based aCSF (in mM) 0.5 CaCl_2_, 7 mM MgSO_4_; NaCl replaced by an equimolar concentration of choline). Slices were incubated for 30min at 32oC in an oxygenated normal aCSF containing (in mM): 126 NaCl, 3.5 KCl, 1.2 NaH_2_PO_4_, 26 NaHCO_3_, 1.3 MgCl_2_, 2.0 CaCl_2_, and 10 D-glucose, pH 7.4, 315 mOsm/l. Slices were allowed to equilibrate for at least 30 min at RT before being transferred to the recording chamber. During recordings, slices were perfused at a rate of 4 ml/min with continuously aerated (95% O_2_-5% CO_2_) normal aCSF at RT.

#### Data Acquisition

Neurons were impaled with patch pipettes (5–7 MΩ) and recorded in the whole-cell configuration, either in the current-clamp or voltage-clamp mode. For current-clamp experiments, the composition of the intracellular solution was: 130 mM K-MeSO_4_, 5 mM KCl, 5 mM NaCl, 10 mM HEPES, 2.5 mM Mg-ATP, and 0.3 mM GTP, 265–275 mOsm, pH 7.3. To measure neuronal excitability and passive membrane properties a 500ms step-pulse protocol was used with current stimulation from −200pA to +50pA. To measure action potential properties, a 5ms step-pulse protocol was used.

For voltage-clamp experiments, the composition of the intracellular solution was: 120 mM Cs-gluconate, 20mM CsCl, 0.1 mM CaCl_2_, 1 mM EGTA, 0.4 mM Na-guanosine triphosphate, 2mM Mg-adenosine triphosphate, 10 mM HEPES. For perforated patch-clamp recordings, patch pipettes were front-filled with the above intracellular solution and back-filled with the same solution containing 5μg/ml gramicidin. No correction from liquid junction potential was applied between the pipette and the aCSF. Whole-cell measurements were low-pass filtered at 5 kHz using an Axopatch 200B amplifier (Molecular Devices, Inc). Recordings were digitized with the ITC-18 board (Instrutech, Inc) on a PC using custom-made codes in IgorPro (Wavemetrics, Inc). All signals were collected at a sampling frequency of 20kHz. For voltage-clamp recordings, the composition of the intracellular solution was:

#### Data Analysis

Data were analyzed using custom-written codes in IgorPro software (Wavemetrics, Inc.). For passive membrane properties, the resting membrane potential (RMP, mV) was measured within 3 min after establishing the whole-cell configuration, and monitored throughout the experiment. To measure input resistance, a 500ms step-pulse protocol was used with current stimulation from −200pA to +50pA. The input resistance (R_in_, MΩ) was measured by plotting the steady-state voltage deflection in an I-V plot and calculating the slope of the best fit line curve (Rin=V/I). The τ_m_ (membrane time constant, ms) was obtained by fitting a single exponential curve to the voltage deflection at −50pA, and the membrane capacitance (C_m_) was calculated using the formula C_m_ = τ_m_/R_in_. In addition, the number of spikes generated in response to a 500ms step-pulse range from +100pA to +300pA was measured.

To measure action potentials (APs) properties, we applied small supra-threshold 5ms step-pulse currents to the cell from −65mV. The active properties were measured at the minimum current stimulation (Rheobase, pA) that generated an AP. The AP threshold (mV) was calculated by taking the first derivative of the voltage trace, defining a threshold and identifying the voltage level at that time point. The rate of rise of the AP (dV/dt, mV/ms) was the maximum value of that first derivative trace. The AP amplitude (mV) was defined as the voltage difference between AP threshold and AP peak. The AP duration (ms) was calculated by the full width of the waveform at the half maximal amplitude (half-width). The fast afterhyperpolarization (fAHP) minimum (mV) was defined as the minimum voltage value right after the AP. The fAHP amplitude (mV) was calculated as the difference between the AHP minimum and the AP threshold. The fAHP time (ms) was defined as the time duration from the time point of AP threshold to the fAHP minimum.

The measurements of spontaneous excitatory postsynaptic currents (sEPSCs) were taken at - 60mV and for spontaneous inhibitory postsynaptic currents (sIPSCs) at +10mV. Automatically selected events were subsequently visually monitored to discard erroneously included noise. The events showing only single peaks were selected for kinetics analysis. All currents detected from every single neuron were averaged. The peak amplitude was calculated as the maximum current value. The time constant of the decay phase was detected by curve fitting with a single exponential decay function.

To investigate the IPSC reversal potential, a stimulating electrode was positioned 100μm away from the recording electrode at the border between layer I and layer II. Recordings were taken at different voltage steps, from −80mV to +40mV in the presence of CNQX (20μM) and AP5 (50μM).

### Immunohistochemistry

Mice at the age of P10 and P20 were perfused with 4% paraformaldehyde, followed by fixation with the same solution for 1h at 4°C, followed by cryoprotection and preparation of 12 μm cryostat sections as previously described (Vidaki et al., 2012). Primary antibodies used were rat monoclonal anti-GFP (Nacalai Tesque, Kyoto, Japan, 1:5000), rabbit polyclonal anti-GFP (1:500; Minotech Biotechnology, Heraklion, Greece) and rabbit polyclonal anti-parvalbumin (PV) (Swant, Bellinzona, Switzerland; 1:2000. Secondary antibodies used were goat anti-rat-Alexa Fluor-488, goat anti-rabbit Alexa Fluor-488, and goat anti-rabbit-Alexa Fluor-555 (Molecular Probes, Eugene, OR, United States, 1:800). Images were obtained with a confocal microscope (Leica TCS SP2, Leica, Nussloch, Germany). For each age group (P10, P20), 2-4 10μm-thick sections from each mouse brain were selected, all including the mPFC and BC.

### RNA *In Situ* Hybridization

Non-radioactive *in situ* hybridization experiments were performed on cryostat sections (12μm thick, see immunochemistry) according to the protocol described (Schaeren-Wiemers and Gerfin-Moser 1993). Riboprobe was prepared by *in vitro* transcription and was specific Somatostatin (SST) (Liodis et al. 2007).

### Nissl Staining

Cryostat sections (12μm thick, see immunochemistry) were incubated in 1:1 100% ethanol:chloroform overnight at RT. Then, sections were rehydrated for 1 min in 100%, 95% ethanol solutions and dH_2_O at RT, followed by a 10-min incubation in 0.1% cresyl violet solution at 50oC. Sections were then dehydrated with dH_2_O, 95%, 100% ethanol and xylene for 5 min and coverslipped with permount. Images from whole sections were obtained in 5× magnification of a light microscope (Axioskop 2FS, Carl Zeiss AG, 268 Oberkochen, Germany) and merged using Adobe Photoshop CC 2015, Adobe Systems, Inc.

### Analysis for Immunochemistry, *in situ* hybridization and Nissl staining

Images taken from Nissl staining slices were analyzed with Matlab, using a custom-made algorithm, which was double checked with hand-counting (Konstantoudaki et al. 2016; Chalkiadaki et, 2019). Images from immunochemistry and in situ hybridization experiments, were hand counted using ImageJ.

### Western blots

Mice were decapitated following cervical dislocation, the brain was quickly removed, placed in ice cold PBS (phosphate-buffered saline) and then positioned on a brain mold, where 1.5 mm slices were taken containing the mPFC and BC. The slices were placed on dry ice, and the prelimbic area of mPFC was dissected out and stored at −80°C. The BC was also isolated from the corresponding slices and stored at −80°C. Frozen tissue blocks were lysed in a solution containing (in mM) HEPES 50, NaCl 150, MgCl2 1.5, EGTA 5, Glycerol 1%, Triton-X100 1%, 1:1000 protease inhibitors cocktail. Proteins ran on 8.5% bis-acrylamide gel and were transferred onto a nitrocellulose membrane (Whatman GmbH, Dassel, Germany). The membrane was blocked, incubated in rabbit polyclonal anti-K+/Cl-Cotransporter (KCC2) (Merck KGaA, Darmstadt, Germany, 1:1000) or rabbit monoclonal anti-GAPDH (Cell Signaling Technology Europe BV, Leiden, Netherlands, 1:1000), washed, incubated in secondary goat anti-rabbit IgG Horseradish Peroxidase Conjugate antibody (Invitrogen, 1:5000), and digitally exposed using the Molecular Imaging system ChemiDoc (BioRad Laboratories, Inc, California, U.S.A.). Analysis of KCC2 and GAPDH expression was performed with ImageJ software, and the raw values of KCC2 from each sample were normalized to their respective GAPDH values.

### Statistical analysis

Statistical analyses were performed in Microsoft Office Excel 2007 and GraphPad Prism Software 7.0. Data are presented as mean ± standard error of mean (SEM). Normality distribution and equality of variances of dataset were tested with the Kolmogorov-Smirnov test normality test. The null hypothesis was rejected for a >5%. When four experimental groups (P10 mPFC, P20 mPFC, P10 BC and P20 BC) were assessed and two variables were taken into consideration (age and brain area), data were analyzed with a two-way ANOVA with Fisher LSD, Sidak’s or Tukey’s multiple comparisons (electrophysiological recordings and cell counting). When three groups (P10 mPFC, P20 mPFC and P10 BC) data were analyzed with one-way ANOVA (electrophysiological recordings). Significant differences were detected by one-way ANOVA. Significance levels of *p < 0.05, **p < 0.01, ***p < 0.001 or ****p < 0.0001 were tested. For comparison of Western blot analysis, the significant effect of each developmental age group from mPFC and BC was assessed using Student’s t-test depending on the experiment.

### Modeling

We adapted the PFC microcircuit model we had developed previously (Konstantoudaki et al., 2014) to fit our experimental data with regards to intrinsic properties of pyramidal neurons and fast-spiking interneurons. We generated two model networks: a) a neonatal mPFC model network and b) a juvenile mPFC model network. In the neonatal mPFC model network the following adjustments were made: a) the pyramidal model neuron was adjusted to fit the passive and active properties of the neurons recorded in this paper, primarily the neonatal pyramidal model neuron exhibited increased input resistance and reduced spike amplitude, b) the fast-spiking interneuron model neuron was adjusted to have reduced fast afterhyperpolarization, as found in our experiments and c) the GABA_A_R receptor model had a reversal potential of −40mV. In the juvenile mPFC model network, the following adjustments were made: a) the GABA_A_R receptor model had a reversal potential of −60mV. All other properties were maintained as in the original model. The network was stimulated with spontaneous synaptic activation on pyramidal model neurons with characteristics similar to the sEPSC and sIPSC properties we recorded. Each condition was simulated in 10 different trials. In each trial, the location of activated synapses varied.

### Data availability

Data presented in the figures in this paper are available upon request. The model networks will be uploaded on ModelDB once published.

## Results

Mice belonging to two age groups were investigated: (i) neonatal mice included pups of postnatal days (P) 9-11 and are defined as P10 while (ii) juvenile mice defined as P20 animals included pups of P19-P21. Due to the high density of intra-cortical synapses in the superficial cortical layers (DeFelipe and Fariñas 1992; Clancy et al. 2001) and their specific involvement in neurodevelopmental disorders (Chini and Hanganu-Opatz, 2020; Bitzenhofer eta l., 2017), we focused on the superficial layers of the mPFC and BC. All analyses that had four groups (mPFC P10 and P20, BC P10 and P20) were conducted using two-way ANOVA, with the two factors being the brain area (mPFC and BC) and age (P10 and P20).

### Synaptic transmission adaptations in mPFC and BC across development

To examine spontaneous synaptic transmission, we performed patch-clamp recordings from layer II/III pyramidal neurons in mPFC and BC from neonatal and juvenile mice. We recorded spontaneous inhibitory postsynaptic currents (sIPSCs, at +10mV) and spontaneous excitatory postsynaptic currents (sEPSCs, at −60mV) and we measured the frequency, amplitude and decay time constant.

In mPFC, the frequency of sIPSCs was significantly augmented in the juvenile group compared to the neonatal group (**Figure 1a,b**), while the sIPSC amplitude and decay-time constant did not significantly change over the investigated time window (**Figure 1a,c,d**). Similarly, the sIPSC frequency and amplitude were significantly increased, in the juvenile compared to the neonatal BC (**Figure 1a,b,c**), while the decay-time constant was not altered (**Figure 1a,d**). On a different note, the sIPSC frequency in BC in the juvenile group was significantly increased compared to the neonatal group in BC but also compared to the juvenile group in mPFC.

**Figure 1.**
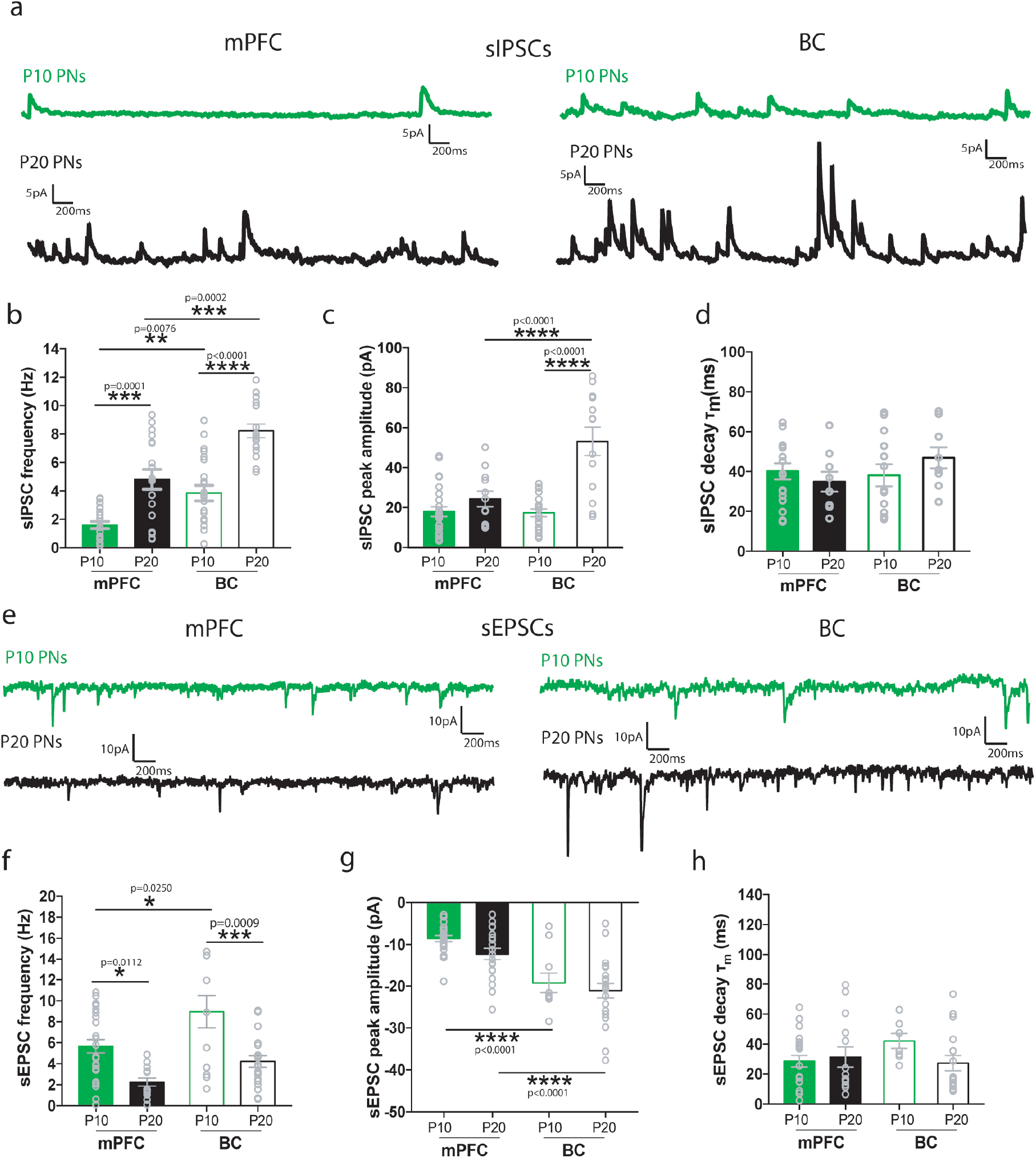
Properties of sIPSCs and sEPSCs at P10 and P20 of layer II/III mPFC and BC pyramidal neurons. **(a)** Representative traces of spontaneous inhibitory postsynaptic currents (sIPSCs) from layer II/III mPFC (left) and BC (right) pyramidal neurons at P10 (green) and P20 (black). **(b)** Bar graph showing the sIPSC frequency (Hz) at P10 and P20 mPFC and BC pyramidal neurons. Two-way ANOVA analyses showed a significant effect of age (F_(1,26)_=19.286, p<0.001) and brain area (F_(1,26)_=12.554, p=0.002) and a trend towards significance of age*brain interaction (F_(1,26)_=3.826, p=0.063). **(c)** Bar graph showing the sIPSC peak amplitude at P10 and P20 of mPFC and BC pyramidal neurons. Two-way ANOVA analyses showed a trend towards significance in the brain area factor (F_(1,26)_=3.420, p=0.077) and no effect of age (F_(1,26)_=1.981 p=0.177) or area*age interaction (F_(1,26)_=0.050, p=0.826). **(d)** Bar graph showing the sIPSC decay time constant (τ_m_) at P10 to P20 of mPFC and BC pyramidal neurons. Two-way ANOVA analyses did not show any significant effect of age (F_(1,26)_=1.129, p=0.299), brain area (F_(1, 26)_=0.211, p=0.651) or area*age interaction (F_(1, 26)_=1.052, p=0.316) was found. **(e)** Representative traces of spontaneous excitatory postsynaptic currents (sEPSCs) from layer II/III mPFC (left) and BC (right) pyramidal neurons at P10 (green) and P20 (black). **(f)** Bar graph showing the sEPSC frequency at P10 to P20 of mPFC and BC pyramidal neurons. Two-way ANOVA analyses showed a significant effect of age (F_(1,25)_=5.273, p=0.032) and brain area (F_(1,25)_=7.388, p=0.013), but not a significant interaction age*brain area (F_(1,25)_=0.591, p=0.450) **(g)** Bar graph showing the sEPSCs peak amplitude at P10 to P20 of mPFC and BC pyramidal neurons. Two-way ANOVA analyses did not reveal significant effect of brain area (F_(1,25)_=0.853, p=0.366), age (F_(1,25)_=0.144, p=0.708) or interaction (F_(1,25)_=0.712, p=0.408). **(h)** Bar graph showing the sEPSCs decay time constant (τm) at P10 to P20 of mPFC and BC pyramidal neurons. Two-way ANOVA analyses showed no significant effect of age (F_(1,25)_=0.557, p=0.463), brain area (F_(1,25)_=0.720, p=0.405) or interaction (F_(1,25)_=1.128, p=0.300). *n=9-13 cells from 5-9 mice/age group.

The sEPSC frequency was significantly decreased in the juvenile compared to the neonatal group, in both areas (**Figure 1e,f**), while the amplitude and decay time constant were unaltered (**Figure 1g,h**). Upon comparing the two brain areas, the sEPSC frequency and amplitude were found significantly decreased in mPFC, compared to BC, in the neonates (**Figure 1f**). In juvenile mice, the sEPSC frequency was similar between the two cortical areas, while the amplitude remained significantly smaller in mPFC compared to BC in both ages (**Figure 1g**). The decay time constant was not different between areas at both ages (**Figure 1h**).

We further investigated the changes in sIPSC and sEPSC frequency changes in the mPFC, in particular, in the presence of AMPA and NMDA receptor antagonists, CNQX and AP5, respectively and a GABA_A_ receptor antagonist, bicuculine. The sIPSC frequency remained increased in the juvenile group, compared to the neonatal group, CNQX and AP5 (**Figure 2a,c**). On the other hand, bicuculine blocked the sIPSCs both in the mPFC at both ages (**Figure 2 b, c**). Application of CNQX eliminated the sEPSCs in juvenile mPFC slices and significantly reduced, but did not eliminate sEPSCs, in neonatal mPFC slices (**Figure 2e,g**). Application of bicuculine (10uM) did not affect the sEPSC frequency in juvenile mPFC slices but significantly reduced sEPSC frequency in the neonatal mPFC (**Figure 2f,g**). These results suggest that depolarizing GABA_A_R currents could contribute to the increased sEPSC frequency observed in layer II/III mPFC pyramidal neurons in the neonatal group.

**Figure 2.**
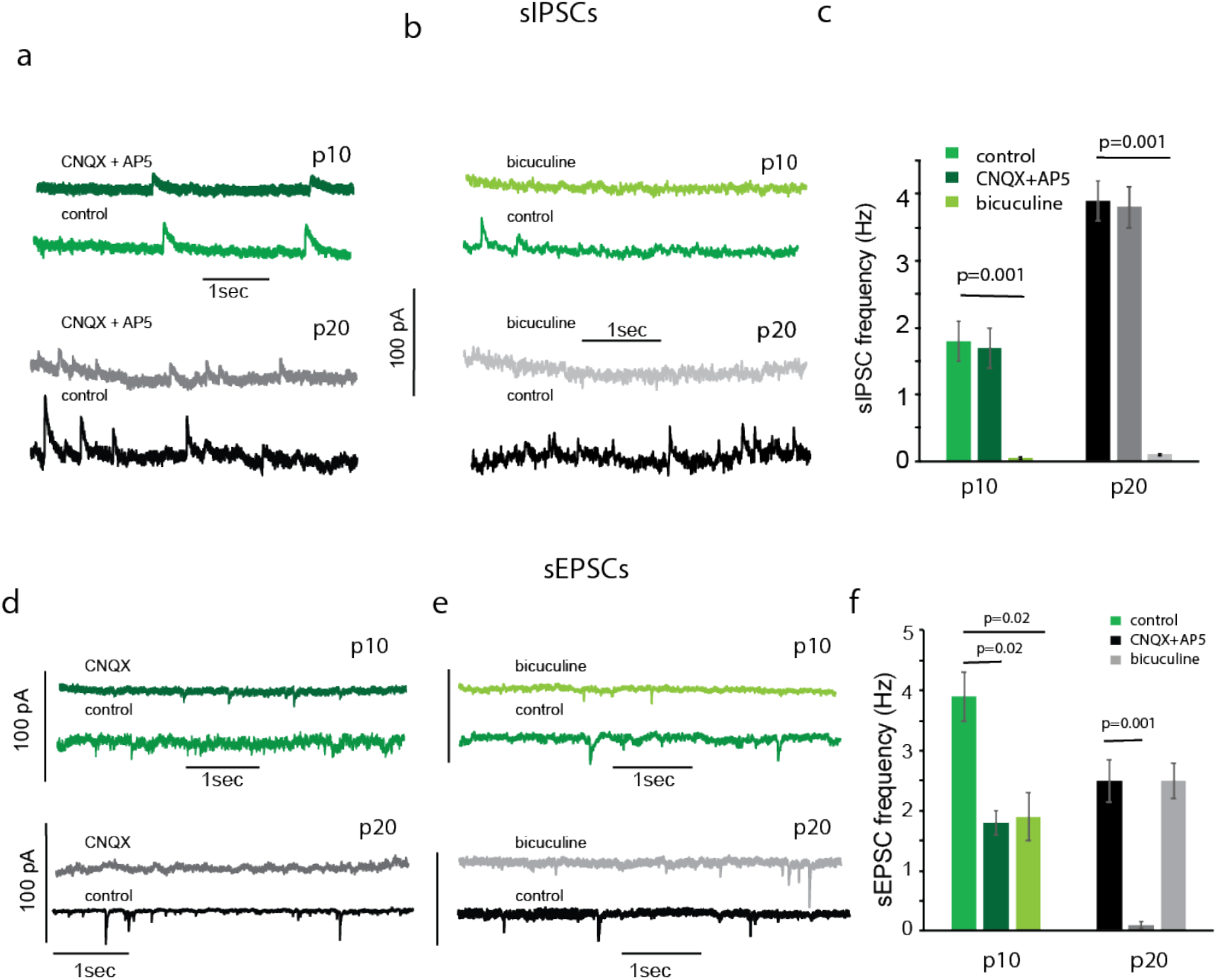
sIPSCs and sEPSCs frequency at P10 and P20 of layer II/III mPFC and BC pyramidal neurons. **(a)** Representative traces showing that CNQX+AP5 did not alter the sIPSC frequency either at P10 or P20 mPFC (recorded at +10mV). **(b)** Representative traces showing that bicuculine blocks sIPSCs either at P10 or P20 mPFC (recorded at +10mV). **(c)** Graphs showing that bicuculine blocks the sIPSCs in P10 and P20 mPFC (t-test, p=0.001), while CNQX+AP5 does not have an effect (t-test, p=0.5). **(d)** Representative traces showing that CNQX reduces sEPSC frequency at p10 and blocks sEPSCs at P20 mPFC (recorded at −70mV). **(e)** Representative traces showing that bicuculine reduces sEPSC frequency at p10 and does not affect sEPSCs at P20 mPFC (recorded at −70mV). **(f)** Graphs showing that at P10 mPFC both CNQX+AP5 and bicuculine reduces the sEPSC frequency in P10 mPFC (t-test, p=0.03 for both drugs), while in P20 mPFC only CNQX+AP5 blocks sEPSCs.

### GABA is depolarizing in the neonatal mPFC but not BC

Our data so far suggests that GABA_A_R function is still depolarizing in the mPFC in neonates. To further investigate this, we performed perforated patch-clamp recordings and recorded evoked IPSCs (eIPSCs) in the mPFC at both ages in the presence of CNQX and AP5 at different voltage steps. A line curve was fit across the eIPSC currents recorded from −60mV to +10mV, because there was a linear V-I relationship at this range. From the graphs, it is evident that reversal potential at P10 is more positive than −30mV, while the reversal potential at P20 is around −45mV (**Figure 3a,b**).

**Figure 3.**
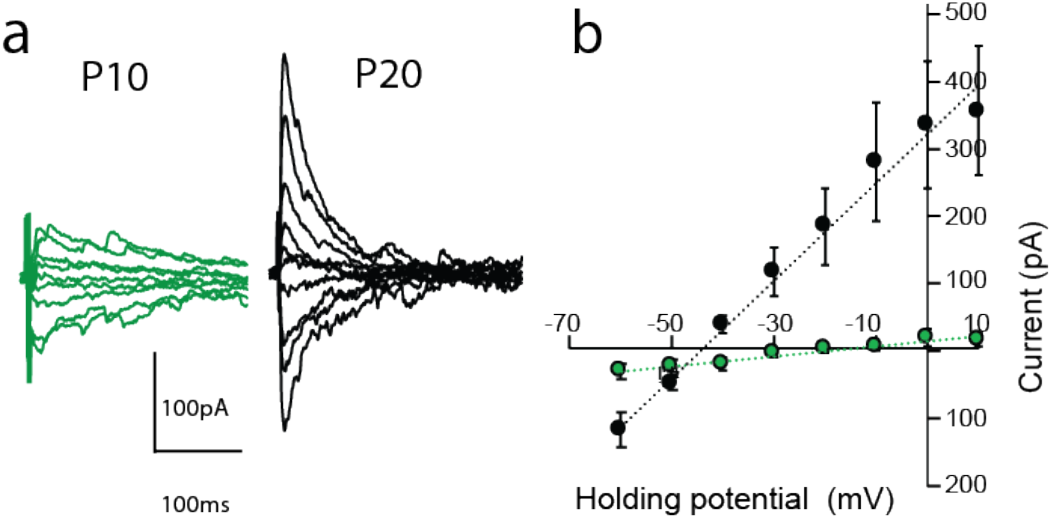
The reversal potential for evoked IPSCs is depolarized at p10 mPFC compared to p20. (a) Representative traces of evoked IPSCs at p10 and p20 mPFC (b) Graph showing the average of eIPSCs at different holding potentials.

We further investigated the GABA_A_R activity by isolating a GABA_A_R response in fEPSP recordings by using CNQX and AP5. The addition of diazepam (2μM) (a GABA_A_R agonist) increased the fEPSP amplitude in mPFC in the neonatal but not juvenile groups (**Figure 4a–b**). In BC, there was no effect of diazepam, at either age (**Supplemental Figure 1**). The switch in the GABA_A_R function from depolarizing to hyperpolarizing occurs due to the increased expression of the K^+^-Cl^−^ co-transporter 2 (KCC2) (Rivera et al. 1999). To determine whether modulating chloride transporters could alter the diazepam-induced enhancement of the fEPSP at P10 mPFC, we recorded the fEPSP in the presence of bumetanide (10uM), which blocks the NKCC1 transporter, and tested the effect of diazepam. We find that in the presence of bumetanide, diazepam did not result in an increase of the fEPSP (**Figure 4c–d**). In addition, we demonstrated that KCC2 protein levels were significantly increased in the juvenile compared to the neonatal mPFC but not in the BC (**Figure 4e–g**). The above results collectively suggest that the GABA_A_R function is depolarizing in the neonatal mPFC.

**Figure 4.**
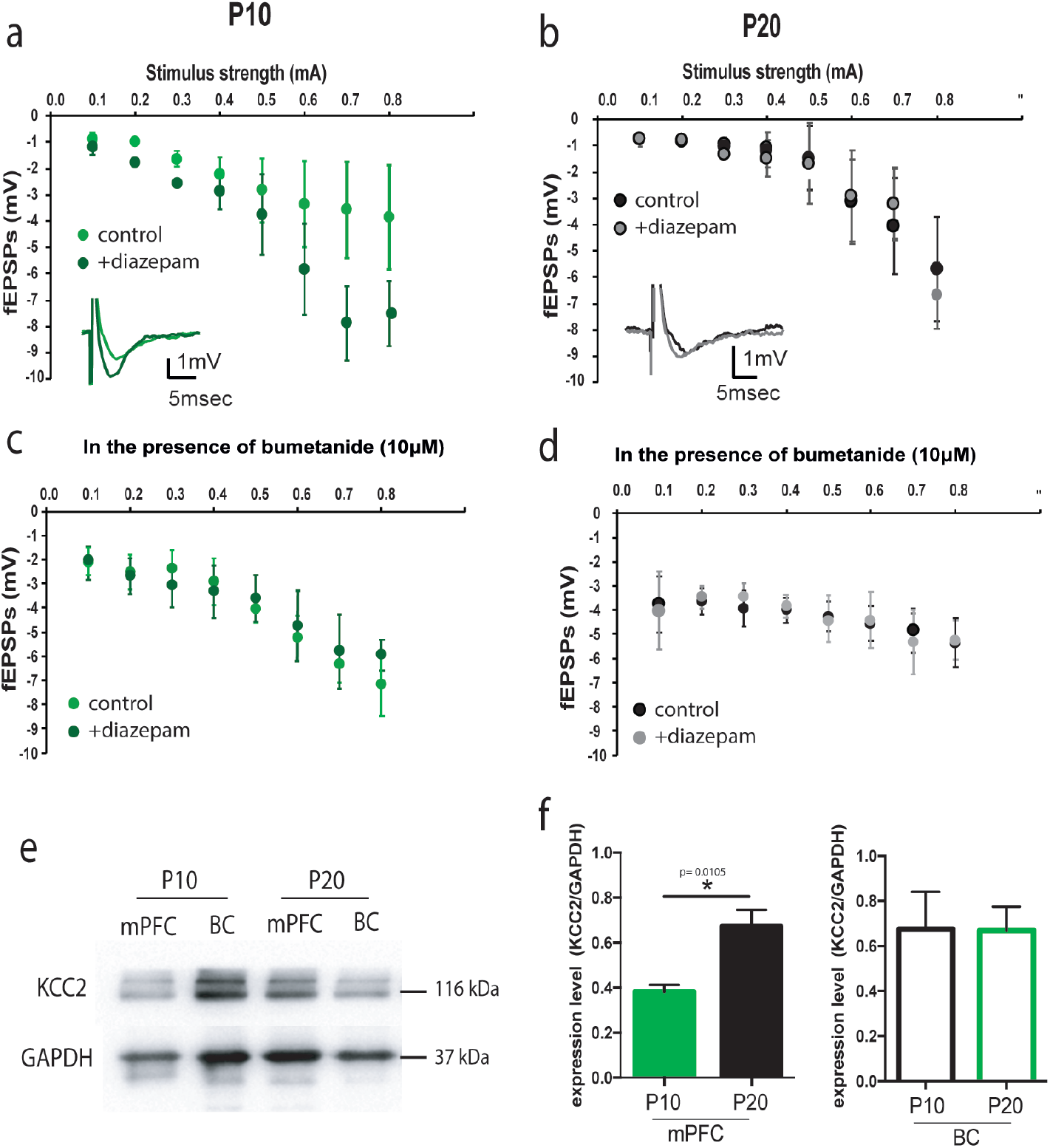
Changes in the chloride transporters mediate the depolarizing action of GABA_A_R at P10 mPFC. fEPSPs were recorded were recorded in layer II/III mPFC in response to current pulses of increasing stimulus strength of layer II/III, during two experimental treatments, before and after application of 2μM diazepam (GABA_A_R agonist) at P10 and P20. **(a)** Representative traces (left) and graph (right) showing the fEPSPs amplitude before (green) and after (dark green) diazepam bath application (in the presence of CNQX and AP5), in mPFC at P10. Two-way repeated measures ANOVA analyses of evoked fEPSPs revealed significant effect of stimulus strength (F_(1,7)_ = 20.64, p=0.004) and experimental treatments (F_(1,7)_ = 5.025, p=0.036) (n=2 brain slices from 3-4 mice). **(b)** Graph (right) and representative traces (left) showing that diazepam bath application does not have any effect on the fEPSP amplitude in mPFC at P20 (in the presence of CNQX and AP5). Two-way repeated measures ANOVA analyses of evoked fEPSPs revealed a significant effect of stimulus strength (F_(1,7)_ = 10.36, p<0.0001) but not experimental treatment (F_(1,7)_=0.03, p=0.9382), (n=2 brain slices from 3-4 mice). **(c)** Graph showing that the effect of diazepam on fEPSP is occluded in the presence of bumetanide (plus CNQX and AP5) at P10. Two-way repeated measures ANOVA analyses of evoked fEPSPs did not reveal a significant effect of experimental treatment (F_(1,7)_=0.08, p=0.752), (n=2 brain slices from 3-4 mice). **(d)** Graph showing the effect of diazepam on fEPSP in the presence of bumetanide (plus CNQX and AP5) at P20. Two-way repeated measures ANOVA analyses of evoked fEPSPs did not reveal a significant effect of experimental treatment (F_(1,7)_=0.06, p=0.831), (n=2 brain slices from 3-4 mice). **(e)** Representative blots showing changes of the K-Cl co-transporter (KCC2) levels, relative to GAPDH at P10 and P20 in mPFC and BC. (f)Graph showing the normalized protein level (KCC2/GAPDH) in mPFC at P10 and P20. The KCC2 protein levels was significantly increased at P20 compared to P10 in mPFC (two-tailed t-test, p= 0.01) but not in BC (two-tailed t-test, p= 0.97) (n=3-4 mice).

### Passive and active membrane properties of MGE-derived interneurons are altered in the mPFC across development

To investigate whether additional changes in interneuron properties are present between neonatal and juvenile groups, we performed current-clamp recordings from layer II/III mPFC and BC of Lhx6^+^ interneurons. For this reason, Lhx6-cre;ROSA26fl-STOP-fl-YFP mice were used in which Lhx6^+^ interneurons express YFP. Lhx6 is expressed by all post-mitotic and mature MGE-derived interneurons (Liodis et al. 2007), therefore, YFP is expressed in MGE-derived interneurons, which include interneurons that express parvalbumin (PV^+^) and somatostatin (SST^+^).

Upon analysis of the passive properties, we found a significant increase in the input resistance and membrane time constant, as well as a significant decrease in the membrane capacitance in the neonatal compared to the juvenile mPFC (**Figure 5**). In addition, the input resistance and the membrane time constant were higher in the neonatal mPFC, compared to BC (at both ages). There was no difference in the resting membrane potential (RMP) between ages and brain areas (**Figure 5, Supplementary Table 1**).

**Figure 5.**
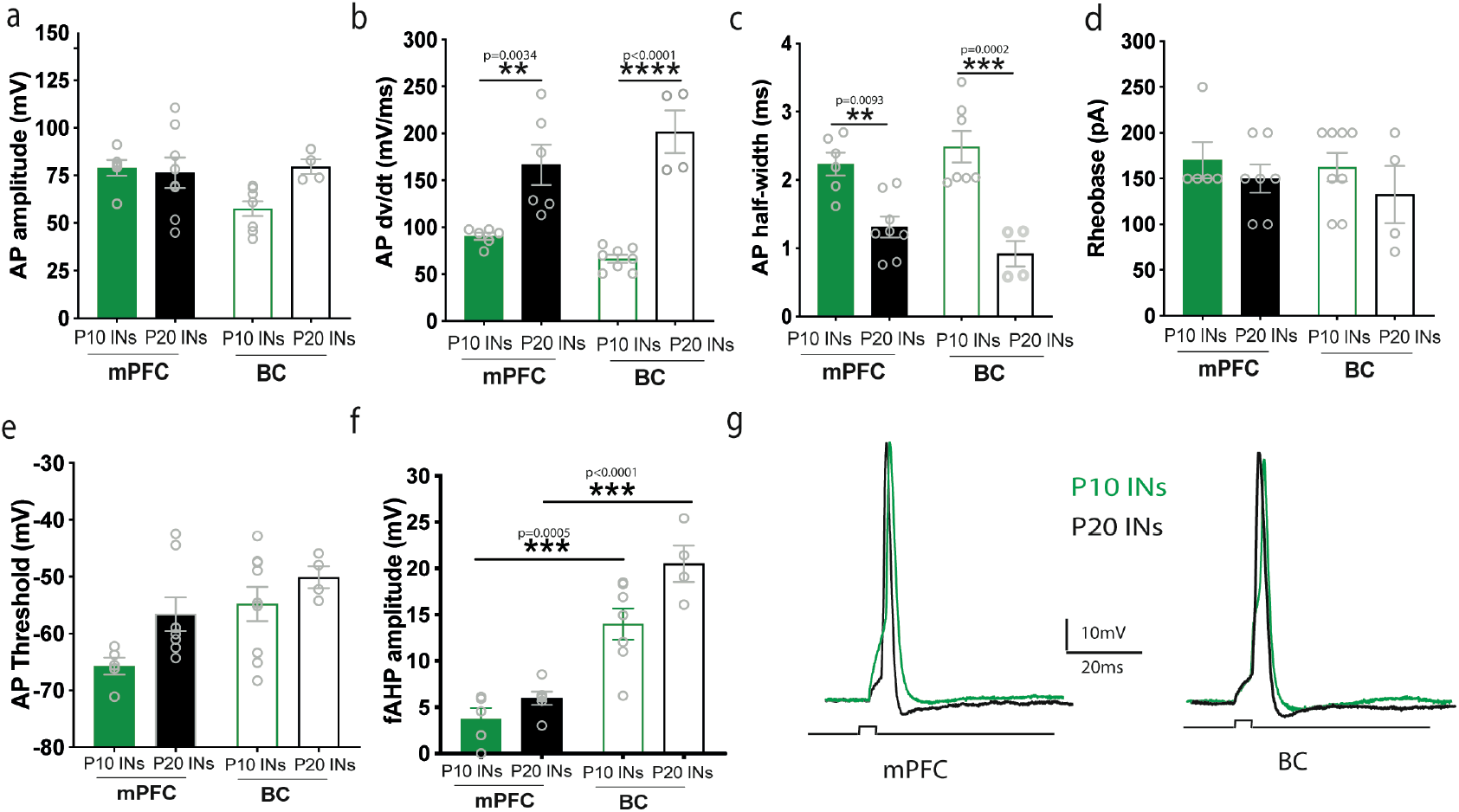
Poor development of active membrane properties of Lhx6^+^ interneurons in mPFC. **(a)** Bar graph showing the action potential (AP) amplitude of interneurons at P10 and P20 in mPFC and BC. Two-way ANOVA analyses did not show any significant effect of age (F_(1,22)_=2.46, p=0.13) or brain area (F_(1, 22)_=2.13, p=0.15) was found., (n=6-9 cells from 5-6 mice/age group). **(b)** Bar graph showing the AP rate of rise (dv/dt) of interneurons at P10 and P20 in mPFC and at P10 in BC. Two-way ANOVA analyses showed a significant effect between ages (F_(1,20)_=58.96, p<0.0001) but not brain area (F_(1,20)_=0.16 p=0.69). Post-hoc analysis showed that the AP rate of rise significantly increased at P20 compared to P10 in mPFC (Tukey’s test, p=0.0034) and at P20 compared to P10 in BC (Tukey’s test, p<0.001), (n=6-9 cells from 5-6 mice/age group). **(c)** Bar graph showing the AP duration (half-width) of interneurons at P10 and P20 in mPFC and BC. Two-way ANOVA analyses showed a significant effect between ages (F_(1,21)_=39.16, p<0.0001) but not brain area (F_(1,21)_=0.16 p=0.73). Post-hoc analysis showed that the AP duration significantly decreased at P20 compared to P10 in mPFC (Tukey’s test, p= 0.0093) and at P20 compared to P10 in BC (Tukey’s test, p=0.0002), (n=6-9 cells from 5-6 mice/age group). **(d)** Bar graph showing the AP rheobase of interneurons at P10 and P20 in mPFC and BC. Two-way ANOVA analyses did not show any significant effect of age (F_(1,20)_=1.60, p=0.22) or brain area (F_(1, 20)_=0.40, p=0.53) was found. **(e)** Bar graph showing the AP threshold of interneurons at P10 and P20 in mPFC and BC. Two-way ANOVA analyses showed significant effect of age (F_(1,22)_=5.048, p=0.035) and brain area (F_(1, 22)_=8.00, p=0.009) was found. Post-hoc analysis showed that the AP threshold was not significantly different at P20 compared to P10 in mPFC (Tukey’s test, p=0.1673) and in BC (Tukey’s test, p=0.72009) or at P10 in mPFC compared to P10 in BC (Tukey’s test, p=0.067) and at P20 in mPFC compared to P20 in BC (Tukey’s test, p=0.72). **(f)** Bar graph showing the AHP (afterhypolarization) amplitude of interneurons at P10 and P20 in mPFC and BC. Two-way ANOVA analyses showed significant effect of age (F_(1,18)_=7.35, p=0.0143) and brain area (F_(1, 18)_=63.72, p<0.0001) was found. Post-hoc analysis showed that the AHP amplitude was not significantly different at P20 compared to P10 in mPFC (Tukey’s test, p=0.7187) and was significantly decreased in mPFC compered to BC, at P10 (Tukey’s test, p= 0.0005) and in mPFC compered to BC at P20 (Tukey’s test, p<0.00001), (n=6-9 cells from 5-6 mice/age group). **(g)** Bar graph showing the AHP time of interneurons at P10 and P20 in mPFC and BC. Two-way ANOVA analyses did not show any significant effect of age (F_(1,19)_=0.009, p=0.92) or brain area (F_(1, 19)_=1.074, p=0.31) was found. **(h)** Representative traces of APs of layer II/III Lhx6+ interneurons in mPFC (left) and BC (right) at P10 (green) and P20 (black).

Regarding the active properties, there was no significant difference between ages and brain areas in the AP amplitude, AP threshold and rheobase, fAHP time (duration) (**Figure 5a, d, e, g; Supplementary Table 1**). The AP rate of rise (dv/dt) was significantly increased while the AP duration (half-width) was significantly reduced in the juvenile compared to the neonatal mPFC and BC (**Figure 6b,c, Supplementary Table 1**). In addition, the fAHP amplitude was significantly lower in the mPFC (**Figure 6f, Supplementary Table 1**), compared to BC. The increased rate of rise and the decreased AP duration are possibly linked with the up-regulation of voltage-dependent sodium channels during development (Huguenard et al. 1988), and in combination with the reduced fAHP amplitude suggest that the MGE-interneurons in neonatal mPFC are still quite immature, when compared to adult PV^+^/SST^+^ interneurons in the same region (Yang et al. 2013; Pan et al. 2017).

**Figure 6.**
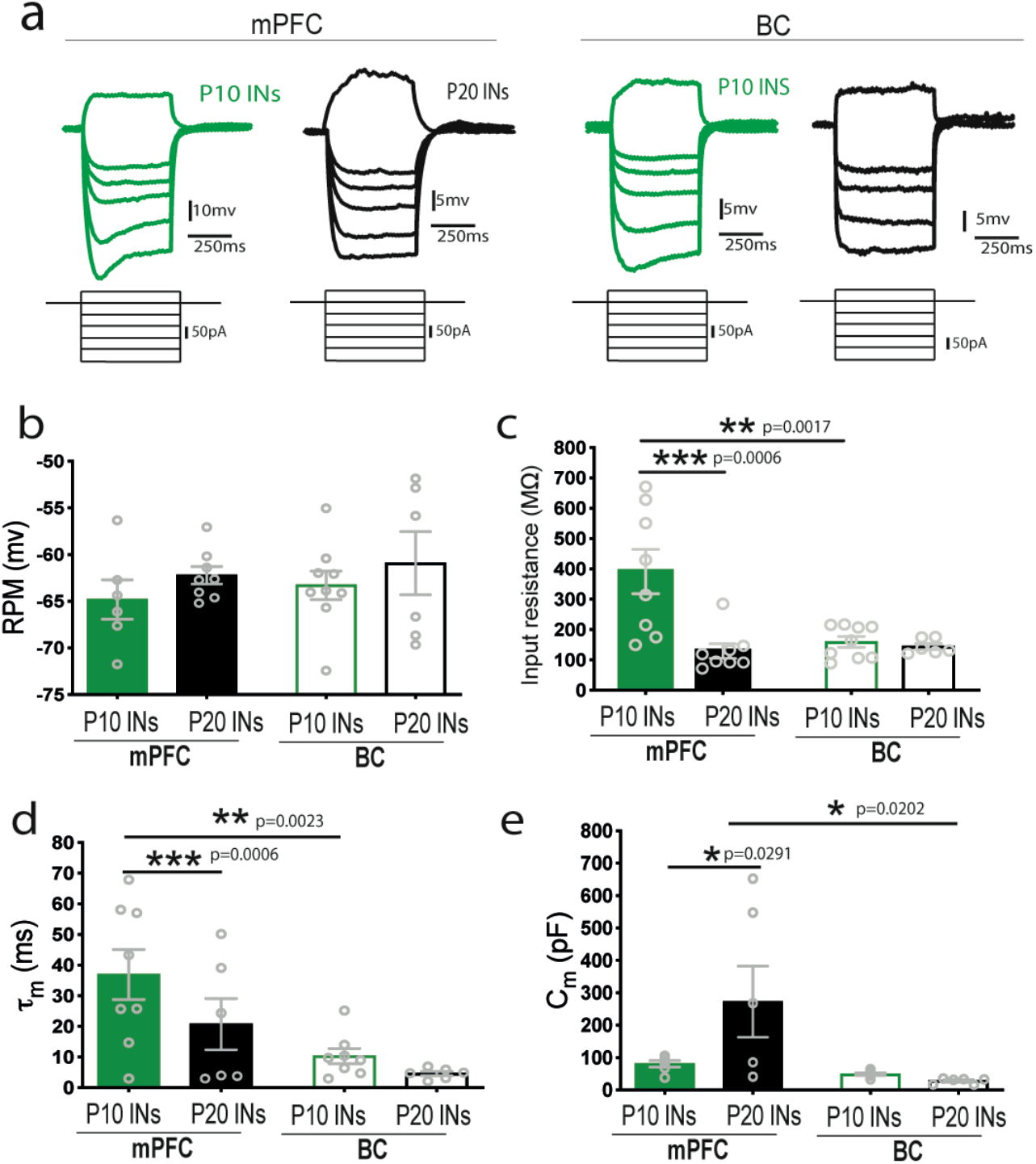
Passive membrane properties of Lhx6+ interneurons at P10 and P20 mPFC and P10 BC. **(a)** Representative voltage responses (top traces) to 500ms positive and negative current pulses (bottom traces, +50, −50, −70, −100, −150, −200 pA) in mPFC at P10 and P20 and BC of Lhx6+ florescent interneurons from layer II/III. **(b)** Bar graph showing the resting membrane potential (RMP) of interneurons at P10 and P20 in mPFC and BC. Two-way ANOVA analyses did not show any significant effect of age (F_(1,25)_=1.55, p=0.22) or brain area (F_(1, 25)_=0.50, p=0.48) was found., (n=6-9 cells from 5-6 mice/age group). **(c)** Bar graph showing the input resistance of interneurons at P10 and P20 in mPFC and BC. Two-way ANOVA analyses showed a significant effect of age (F_(1,27)_=10.94, p=0.0027) and brain area (F_(1,27)_=6.65 p=0.0157). Post-hoc analysis showed that the input resistance significantly decreased at P20 compared to P10 in mPFC (Tukey’s test, p=0.0006) and was significantly higher in mPFC compared with BC, at P10 (Tukey’s test, p=0.0017), (n=8-9 cells from 5-6 mice/age group). **(d)** Bar graph showing the membrane time constant (τ_m_) of interneurons at P10 and P20 in mPFC and BC. Two-way ANOVA analyses showed a significant effect of age (F_(1,24)_=14.71, p=0.0008) and brain area (F_(1,24)_=6.92 p=0.0147). Post-hoc analysis showed that τ_m_ was significantly higher at P10 compared to P20 in mPFC (Tukey’s test, p=0.0006) while it was significantly higher in mPFC compared to BC, at P10 (Tukey’s test, p=0.0023), (n=8-9 cells from 5-6 mice/age group). **(e)** Bar graph showing the membrane capacitance (C_m_) of interneurons at P10 and P20 in mPFC and BC. Two-way ANOVA analyses showed a significant effect between brain areas (F_(1,21)_=6.82, p=0.00163) and not between ages (F_(1,21)_=2.60 p=0.1219). Post-hoc analysis showed that Cm was significantly higher at P10 compared with P20 in mPFC (Tukey’s test, p=0.0291) and was not significantly different between mPFC and BC, at P10 (Tukey test, p=0.97) while it was significantly higher at P20 in mPFC compared to P20 in BC (Tukey test, p=0.0202), (n=6-9 cells from 5-6 mice/age group).

Overall, our data indicate that some intrinsic properties of interneurons in mPFC change with age (from P10 to P20), reaching values that closer resemble adult MGE-derived interneurons (Yang et al. 2013; Pan et al. 2017). The increased sIPSC frequency of mPFC pyramidal neurons observed in the juvenile group could partly be explained by these altered properties of presynaptic interneurons.

### Decreased PV interneurons in mPFC compared to BC

An additional explanation for the adaptations in inhibitory transmission could come from alterations in interneuron cell densities. To test this, we quantified the number of interneurons per area in cryosections in the mPFC and BC at both ages, in coronal brain slices of Lhx6^+^-expressing mice. The YFP^+^ positive cells per area (i.e. Lhx6^+^ cell density) in mPFC and BC was similar between ages, but was significantly lower in the mPFC, compared to BC (**Figure 7a**). However, we found no contribution of cell death in the mPFC interneuron population, as the percentage of cell death in Lhx6^+^ neurons is very low in both mPFC and BC in the neonatal group (**Supplementary Figure 2**).

**Figure 7.**
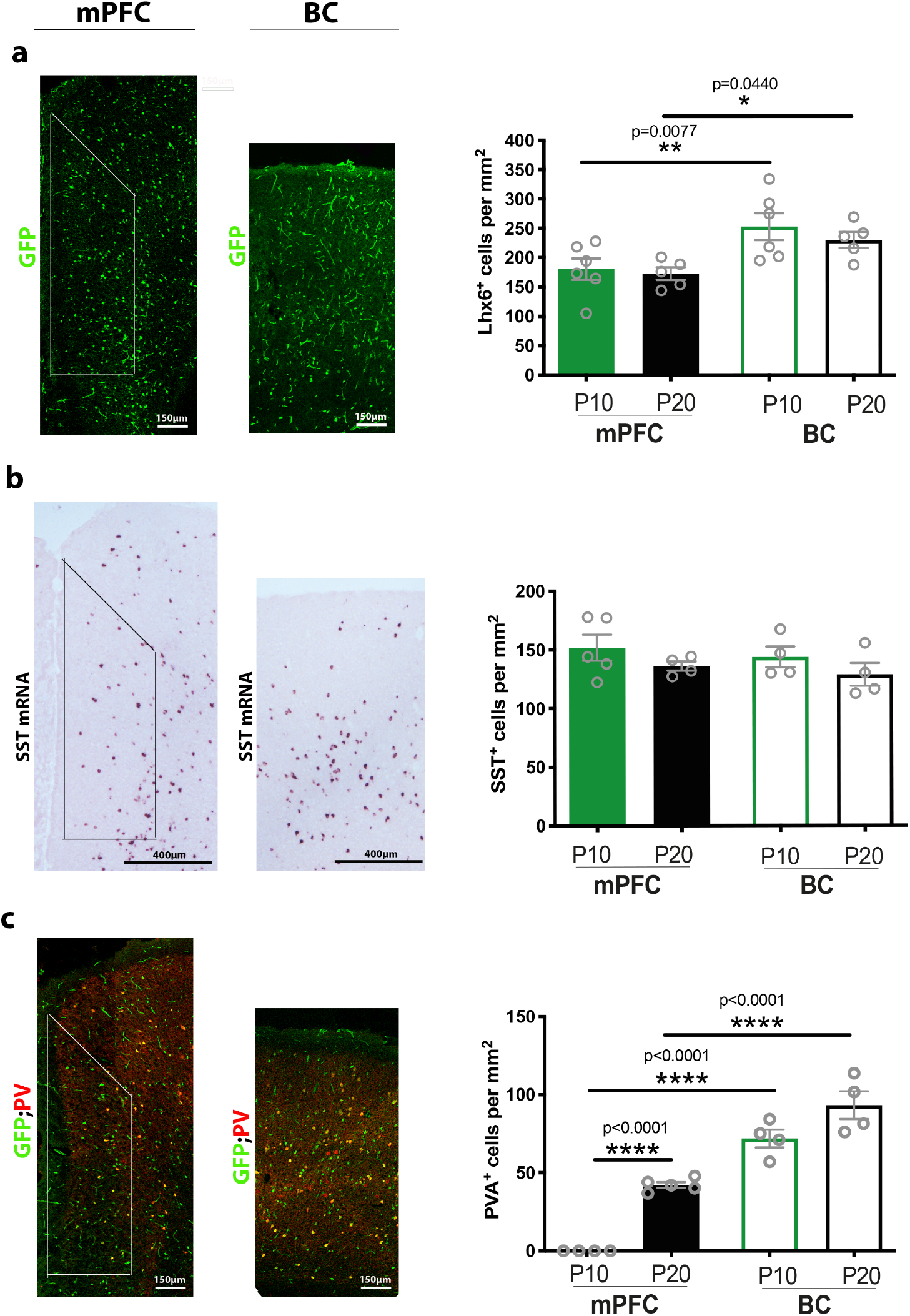
Significant differences in cellular density of Lhx6^+^ interneurons in mPFC and BC at P10 and P20. **(a)** A representative immunostaining with GFP for Lhx6^+^ interneurons in Lhx6-cre;ROSA26fl-STOPfl-YFP mice in mPFC and BC at P20 is showing on the left. Scale bars: 150 μm. On the right, bar graph comparing Lhx6^+^ interneurons cell density (per mm^2^) at P10 and P20 in mPFC and BC. Two-way ANOVA analyses of the cell density revealed a significant effect of brain area (F_(1, 18)_ = 13.11, p=0.0020), but not of age (F_(1,18)_=0.7185, p=0.4078). Post-hoc analysis showed that the Lhx6^+^ cell density was not significant different at P20 compared to P10 in mPFC and BC (LSD test, p=0.77 and p= 0.38, respectively). The Lhx6^+^ cell density was significantly lower in mPFC compared to BC at P10 and P20, respectively (LSD test, p=0.0077 and p= 0.0440, respectively), (P10 in mPFC and BC:n= 5 mice, P20 in mPFC and BC: n=4). **(b)** A representative in situ hybridization staining for somatostatin positive cells (SST^+^) using wild type animals in mPFC and BC at P20 is showing on the left. Scale bar: 200μm. Bar graph comparing cell density based on SST^+^ expression at P10 and P20 in mPFC and BC. Two-way ANOVA analyses of the cell density showed no significant effect of age (F_(1, 13)_= 2.78, p=0.12) and brain area (F_(1,13)_=071, p=0.41) was found, (P10 in mPFC: n= 5 mice, P10 in BC and P20 in mPFC and BC: n=4). **(c)** A representative double immunostaining for GFP; PV (PV: parvalbumin) in mPFC and BC at P20 is showing on the left. Scale bars: 150 μm. On the right, bar graph comparing cell density based on PV^+^ expression at P10 and P20 in mPFC and BC. Two-way ANOVA analyses of the cell density revealed a significant effect of age (F_(1, 14)_ = 45.49, p<0.0001) and brain area (F_(1, 14)_ = 170.2, p<0.0001). PV^+^ cells were not found in mPFC but were identified in BC, at P10. Post-hoc analysis showed that the PV^+^ cell density was not significantly different at P20 compared to P10 in BC (LSD test, p= 0.1089), but was significantly lower in mPFC compared to BC at P20 (LSD test, p<0.0001), (P10 and P20 in mPFC: n= 5 mice, P10 and P20 in BC: n=4).

The transcription factor Lhx6 is required for the specification and maintenance of main MGE-derived interneurons, PV and SST-positive interneuron subtypes, at postnatal ages (Liodis et al. 2007). The neuropeptide SST (both mRNA and protein) is progressively expressed from embryonic to postnatal levels (Bendotti et al. 1990; Forloni et al. 1990). We found that the SST mRNA levels were similar between areas and ages (**Figure 7b**). On the other hand, the emergence of PV immunoreactivity in the mouse cortex shows a delayed development, starting from early postnatal period to adult, with marked area-specific differences(Del Rio et al. 1992). We found that PV was only immunoreactive in BC, and not in mPFC, at P10 (**Figure 7c and Supplementary Figure 3**). At P20, PV was immunoreactive in both mPFC and BC, but PV^+^ cell density was significantly lower in the mPFC, compared to BC (**Figure 7c**).

We also counted the total cell density of mPFC and BC from neonatal and juvenile mice (**Supplementary Figure 4a**) using Nissl staining. In the mPFC, the cell density significantly decreased at P20 compared to P10 (**Supplementary Figure 4b**). On the contrary, in BC, the total cell density significantly increased at P20 compared to P10 (**Supplementary Figure 4b**). When the two brain areas were compared, no difference was found at P10, while the mPFC cell density was significantly lower compared to BC at P20 (**Supplementary Figure 4b**).

We further examined whether the alterations in total cell density are derived from alterations in cell density of interneurons by measuring the Lhx6^+^ neurons over the Nissl-positive cells. No differences were detected between areas and ages (**Supplementary Figure 4c).** These results suggest that the changes in total cell density in mPFC and BC respectively are probably due to changes in other neuronal or glial populations.

### No significant changes in pyramidal neuron excitability between the two brain regions

To determine whether the reduced sEPSC frequency can be explained by changes in pyramidal neuron excitability, we investigated their intrinsic properties. The passive and active properties of pyramidal neurons were measured using current-clamp recordings from layer II/III mPFC and BC. With regards to passive properties, no significant differences were observed in the RMP, the input resistance and the membrane time constant between brain regions and ages (**Supplementary Figure 5, Supplementary Table 1**). Only the membrane capacitance was significantly increased at P20 compared to P10 (**Supplementary Figure 5d, Supplementary Table 1**), in both brain areas. In addition, the number of spikes generated with increasing current stimulation was not significantly different between ages and regions (**Supplementary Figure 6**). In terms of active properties, the AP amplitude and rate of rise were increased at P20 compared to P10 mPFC, while the AP half-width, rheobase and threshold were not significantly different (**Figure 8, Supplementary Table 1**). The AP amplitude was also significantly increased at P20, compared to P10 in BC, while the other properties did not change (**Figure 8, Supplementary Table 1**). Comparing the two regions at the two ages, we found no significant differences of AP properties of pyramidal neurons (**Figure 8, Supplementary Table 1**). The developmental increase of AP amplitude and rate of rise in the mPFC could be due to the on-going maturation of sodium channels in pyramidal neurons. However, these changes could not account for the reduced sEPSCs in the neonatal, compared to juvenile, mPFC and BC.

**Figure 8.**
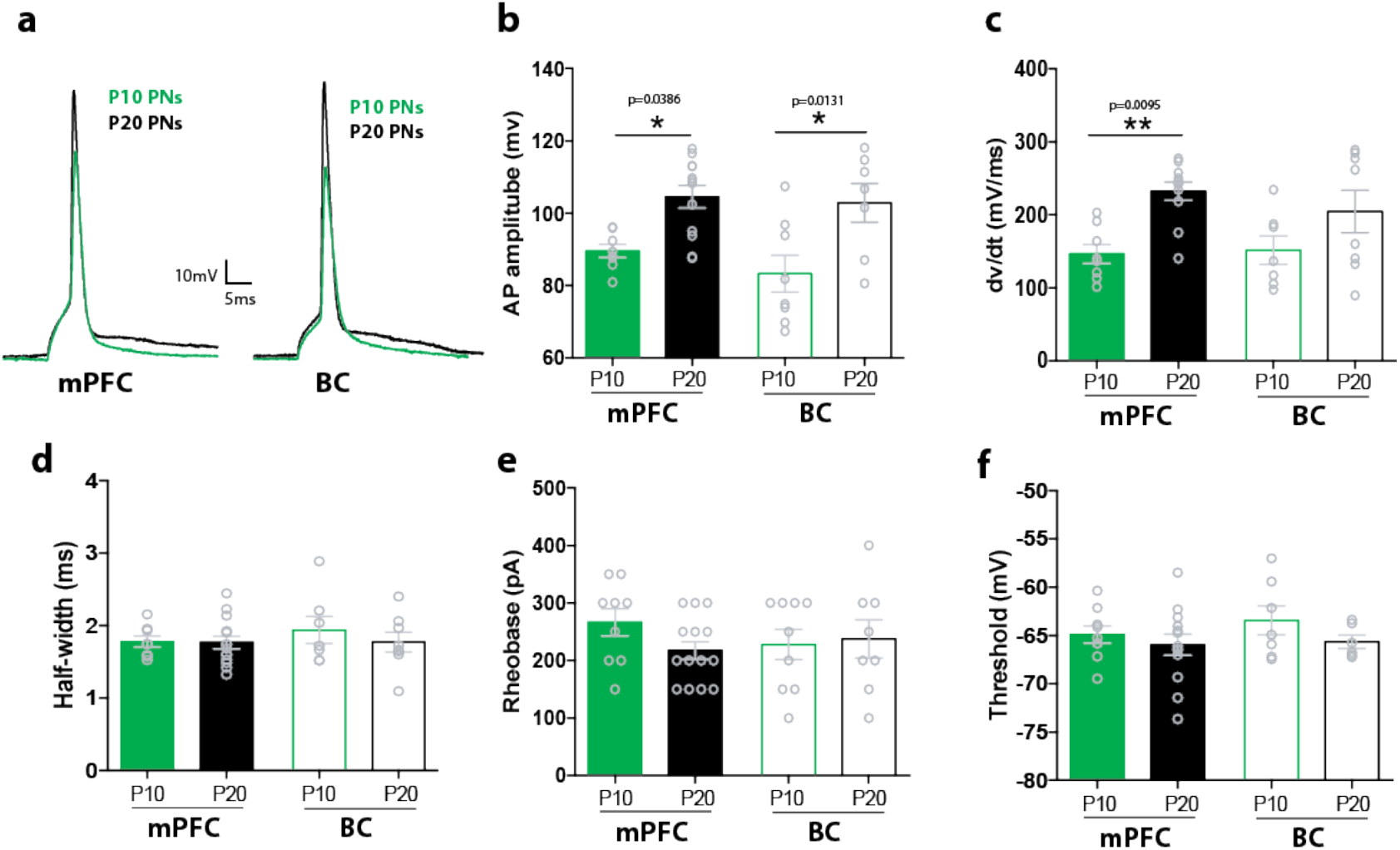
Active properties of mPFC and BC pyramidal neurons. **(a)** Representative traces of action potentials (APs) of layer II/III pyramidal neurons in mPFC (left) and BC (right) at P10 (green) and P20 (black), respectively. **(b)** Bar graph showing the AP amplitude of pyramidal neurons at P10 and P20 in mPFC and BC. Two-way ANOVA analyses showed a significant effect of age (F_(1, 31)_ = 18.74, p=0.0001) but not on brain area (F_(1, 31)_ = 0.99, p=0.32) was found. Post-hoc analysis showed that the AP amplitude significantly increased at P20 compared to P10 in mPFC and BC (Tukey’s test, p=0.0386 and p= 0.0131, respectively) (n=9-14 cells from 6-10 mice/age group). **(c)** Bar graph showing the AP rate of rise (dv/dt) of pyramidal neuron at P10 and P20 in mPFC and BC. Two-way ANOVA analyses showed a significant effect of age (F_(1,30)_= 13.53, p=0.0009) but not on brain area (F_(1, 30)_ = 0.36, p=0.55) was found. Post-hoc analysis showed that the AP rate of rise significantly increased at P20 compared to P10 in mPFC (Tukey’s test, p= 0.0095), but not in BC (Tukey’s test, p= 0.25) (n=8-14 cells from 6-10 mice/age group). **(d)** Bar graph showing the AP duration (half-width) of pyramidal neuron at P10 and P20 in mPFC and BC. Two-way ANOVA analyses showed no significant effect of age (F_(1, 33)_ = 0.52, p=0.47) or brain area (F_(1, 33)_ = 0.43, p=0.51) was found (n=9-14 cells from 6-10 mice/age group). **(e)** Bar graph showing the AP rheobase of pyramidal neuron at P10 and P20 in mPFC and BC. Two-way ANOVA analyses showed no significant effect of age (F_(1, 36)_ = 0.66, p=0.41) or brain area (F_(1, 36)_ = 0.16, p=0.69) was found (n=9-14 cells from 6-10 mice/age group). **(f)** Bar graph showing the AP threshold of pyramidal neuron at P10 and P20 in mPFC and BC. Two-way ANOVA analyses showed no significant effect of age (F_(1,31)_ = 1.90, p=0.17) or brain area (F_(1,31)_=0.55, p=0.46) was found (n=9-14 cells from 6-10 mice/age group).

### Effects of mPFC changes in intrinsic and synaptic properties on PFC network activity

In an effort to understand the circuit effects of the aforementioned differences in intrinsic and synaptic properties between neonatal and juvenile mPFC, we adapted an already validated PFC network model (Konstantoudaki et al., 2014) to our current data to generate a neonatal (P10) mPFC model network and juvenile (P20) mPFC model network. Both model networks were stimulated with spontaneous excitatory and inhibitory inputs based on our results for sEPSC and sIPSC frequency and amplitude at both ages (**Figure 1 and Supplementary Table 1**). Our simulation results predict that the intrinsic and synaptic changes from P10 to P20 in mPFC result in enhanced mPFC network activity at P20, compared to P10 (**Figure 9a,e**), as indicated by single-cell activity (**Figure 9a,b**) and the filtered signal (**Figure 9c,d**). Furthermore, we investigated which adaptations have a significant contribution to the network activity properties observed in each model network. Therefore, we generated neonatal (P10) model networks in which the pyramidal model neuron used had P20 properties (N2 network), the fast-spiking interneuron model used had P20 properties (N3 network) or the GABA_A_R reversal potential was that of P20 (N4 network). We observed that in all the above networks, the resulting neuronal activity generated was significantly reduced compared to the control P10 network (N1) (**Figure 9f**), suggesting that the ‘immature’ properties of pyramidal neurons, interneurons and GABA_A_ receptors allow the neonatal mPFC network to maintain spontaneous neural activity. Similarly, we generated juvenile model networks in which the pyramidal model neuron used had P10 properties (J2 network), the fast-spiking interneuron model used had P10 properties (J3 network) and the GABA_A_R reversal potential was that of P10 (J4 network). We find that the resulting neuronal activity generated was significantly increased compared to the control P20 network (J1) (**Figure 9g**). These data support the notion that the maturation of pyramidal neurons and the GABAergic system contributes to restricting network activity in the juvenile mPFC.

**Figure 9.**
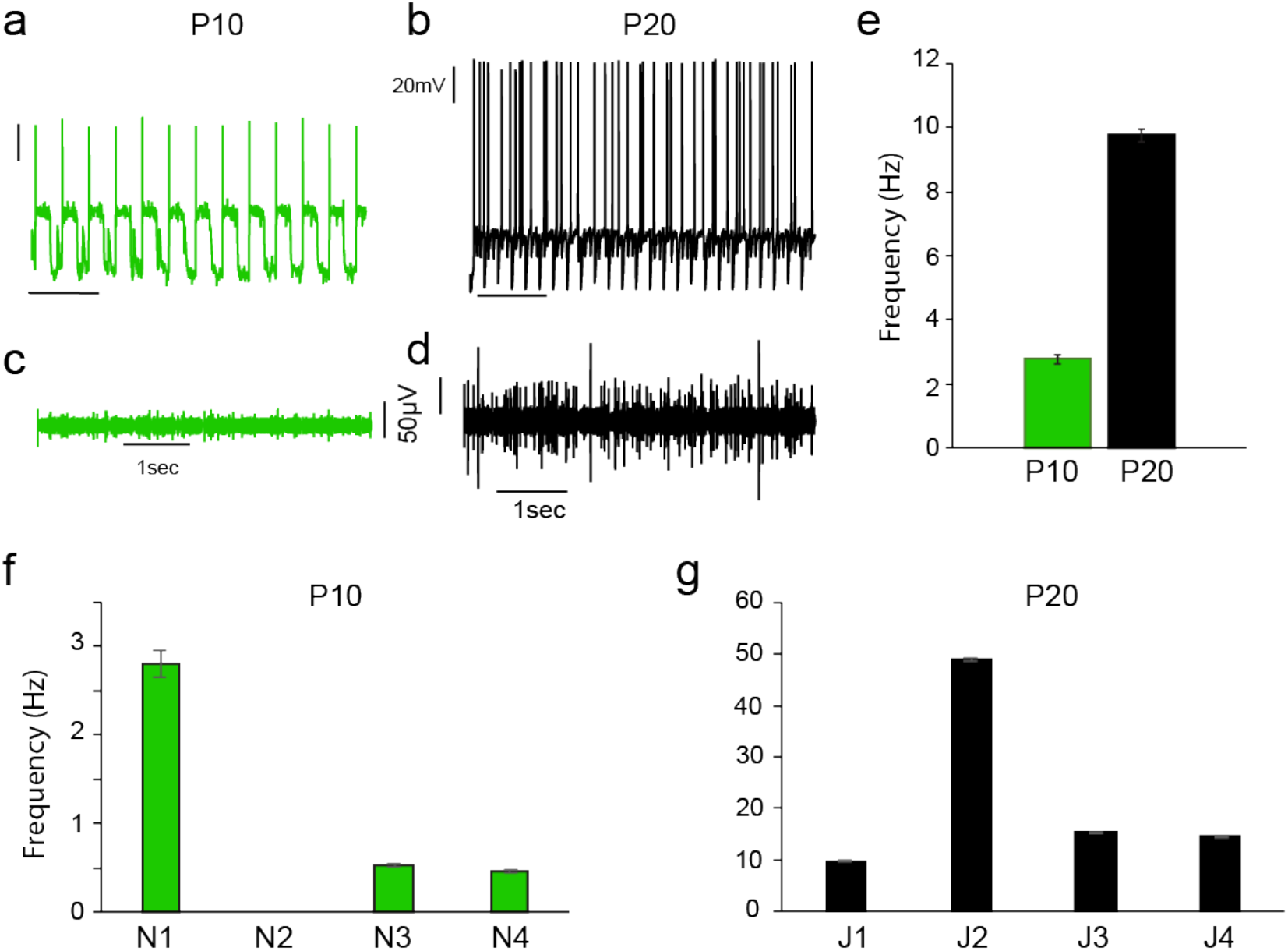
Modeling p10 and p20 PFC network activity. ***(a,b)*** Neuronal response of a single neuron within the PFC model network at p10 and p20 ***(c, d)*** The filtered voltage response of the p10 and p20 model networks **(e)** Average frequency of the P10 and P20 model networks. **(f)** Graph showing how the responses change in modified P10 model networks. N1: control P10 model network, N2: pyramidal model neuron of P20 model network used, N3: fast-spiking interneuron model network of P20 model network used, N4: GABAAR reversal potential was set to −60mV (that of P20 model network). **(g)** Graph showing how the responses change in modified P20 model networks. J1: control P20 model network, J2: pyramidal model neuron of P10 model network used, J3: fast-spiking interneuron model network of P10 model network used, J4: GABAAR reversal potential was set to −40mV (that of P10 model network).

## Discussion

Our study has identified significant developmental events in the mPFC and the BC between the second and third postnatal weeks. Specifically, we have shown that spontaneous inhibitory transmission, measured as sIPSCs, is increased in mPFC from P10 (neonatal) to P20 (juvenile group). Moreover, our data support a depolarizing action of GABA_A_R in the second postnatal week, in the mPFC, as indicated by the presence of non-AMPAR-mediated sEPSCs, depolarizing reversal potential of evoked GABA_A_R responses, increased basal fEPSPs following GABA_A_R activation, which is blocked by concurrent bumetanide application, and decreased protein levels of KCC2. In parallel, differences the intrinsic properties of GABAergic interneurons of mice in the juvenile period resemble are more mature, compared to the neonatal period in mPFC. The above developmental adaptations, along with increased AP amplitude and AP rate-of-rise in pyramidal cells age result in augmented network activity in the juvenile, compared to the neonatal mPFC.

### Depolarizing action of GABA in the immature cortex

GABA plays a crucial role in inhibiting adult neurons, acting primarily via the chloride-permeable GABA_A_R and resulting in hyperpolarization of the membrane potential (Kaila and Voipio 1987). However, GABA action leads to depolarization of immature neurons (i.e. during the first postnatal week in mice), due to an initially higher intracellular chloride concentration [Cl^−^]_in_ (Ben-Ari 2001; Ben-Ari et al. 2007; Ben-Ari 2012). The developmental switch of GABA action from depolarizing to hyperpolarizing results from changes in cation-chloride co-transporter expression: NKCC1, a cation-Cl^−^ importer, is highly expressed in neuronal precursor cells during early brain development (Plotkin et al. 1997; Yamada et al. 2004), while the expression of the K^+^-Cl^−^ cotransporter 2 (KCC2), a cation-Cl^−^ exporter, increases after the first postnatal week (Ben-Ari 2001; Ben-Ari et al. 2007; Ben-Ari 2012). This increased KCC2 transporter expression might provide a central mechanism for the depolarization to hyperpolarization switch of GABAergic transmission via progressive reduction of [Cl^−^]_in_ (Lu et al. 1999; Rivera et al. 1999; Ganguly et al. 2001; Ben-Ari 2002; Dzhala et al. 2005; Fiumelli et al. 2005).

The GABA_A_R switch from depolarizing to hyperpolarizing occurs at P7 in the hippocampus, cortex, amygdala (Ben-Ari et al. 1989; Luhmann and Prince 1991; LoTurco et al. 1995; Owens et al. 1996; Martina et al. 2001; Gulledge and Stuart 2003; Ben-Ari et al. 2007). In the mPFC, GABA application via an iontophoretic pipette and perforated recordings from pyramidal neurons revealed that the reversal potential of GABAA IPSCs is significantly depolarized at P10 and switches around P14 for dendritic IPSCs and after P20 for axon initial segment IPSCs (Rinetti-Viargas et al., 2017). Our results reinforce the GABA_A_ polarity changes after P10 using perforated IPSC recordings following electrical stimulation in layer II mPFC pyramidal neurons and further supported by decreased levels of KCC2 transporter in the neonatal mPFC. Our results contribute to the more detailed understanding of the protracted maturation of mPFC compared to other cortical areas.

### Interneurons and mPFC development

Besides the GABAA receptor properties, the number and electrophysiological properties of interneurons change during early development. Our knowledge on the GABAergic interneurons of neonatal mPFC is very limited. PV expression is lowest in juveniles and increases during adolescence to levels similar to those observed in adulthood (Caballero et al. 2014). Furthermore, PV expression is not evident in the neonatal period and emerges during the juvenile period in the mPFC (del Rio et al. 1994; de Lecea et al. 1995; Zheng et al. 2011; Spampanato and Sullivan 2016). Our results agree with these findings, as PV expression was not detected neonatally and was detected in very low amounts during the juvenile period in the mPFC. On the other hand, SST mRNA levels were not altered between the neonatal and juvenile period in the mPFC or the somatosensory cortex.

Recordings of Lhx6^+^-interneurons indicate that both passive and active properties are regulated by age and reach values that better resemble adult MGE-derived interneurons. Specifically, we have found that the input resistance and AP width decrease while the AP rate of rise increases in the mPFC at P20 compared to P10. In part, similar findings have been identified for PV^+^ cells in the hippocampus (Doischer et al. 2008; Miyamae et al. 2017) and SST^+^ cells in the anterior cingulate cortex (Pan et al. 2017). On the other hand, the AHP amplitude is still quite immature in the mPFC at P20, compared to PV^+^, SST^+^ interneurons in primary sensory areas or the hippocampus and compared to adult mPFC (Doischer et al. 2008; Yang et al. 2013; Pan et al. 2017). Therefore, it is likely that the physiological properties of PV^+^ and SST^+^ interneurons in the mPFC continue to change past the third postnatal week.

The more mature intrinsic electrophysiological properties of GABAergic interneurons likely contribute to the identified increased frequency of sIPSCs between the second and third postnatal weeks. These results are consistent with the developmental changes of IPSCs in layer III pyramidal neurons of monkey PFC (González-Burgos et al. 2015) and mouse mPFC (Kroon et al. 2019).

### Pyramidal neurons and network activity

It has been suggested that spontaneous network activity changes from local, highly synchronized to more diffuse from the second to the third postnatal weeks, in the primary sensory cortices (Golshani et al. 2009; Frye and MacLean 2016). In the mPFC, increased levels of network activity have been recorded from P11 to P23 (Bitzenhofer et al., 2021). Our modeling results, which are based on the electrophysiological properties of pyramidal neurons and interneurons we recorded in this study, all reproduce the increased network activity from the neonatal to juvenile ages. A significant contribution to the changes in network activity comes from the changes observed in pyramidal neurons, in particular, the developmental increase of AP amplitude and rate of rise, which could be due to the on-going maturation of sodium channels in pyramidal neurons.

Studies in developing mPFC pyramidal neurons have proposed that there is a unique sensitive time window for synaptic maturation of these neurons from individual cortical layers. During rat mPFC layer V development, the intrinsic properties, synaptic inputs and morphology of pyramidal neurons develop together during early postnatal life. While the greatest changes were reported during the first ten days after birth, the adult-like properties emerged after the end of the third week (P21) (Zhang et al. 2011). This study confirms that the second postnatal week is a period of rapid growth, similar to that in other neocortical regions by combining functional and structural measurements of developing pyramidal neurons in mouse mPFC (Zhu 2000; Romand et al. 2011).

### Developmental PFC malformations lead to cognitive disorders in adulthood

The neonatal functional maturation of GABAergic circuits and E/I (excitation to inhibition) balance are critical for PFC-dependent behaviours and plasticity in the adult while their malfunction leads to many psychiatric disorders (Benes 1991; Kilb 2012; Ferguson and Gao 2018). From the prenatal period to late adolescence, the PFC network is highly vulnerable to genetic and environmental factors (Andersen 2003), since the mPFC is one of the latest cortical regions to develop (Huttenlocher 1990). While many studies have focused on understanding several developmental processes during adolescence (Caballero et al. 2016), our knowledge regarding the ongoing cellular and network developmental processes during the perinatal period is notably limited, despite significant evidence showing that environmental manipulations during this period manifest as complex psychiatric and neurologic disorders in adulthood (Weinberger 1986).

The delayed developmental shift of GABA action in various mouse models mimicking human brain disorders have been investigated, including the maternal immune activation model (Corradini et al. 2018; Fernandez et al. 2018), the Scn1a and Scn1b mouse models of Dravet syndrome (Yuan et al., 2019), the 22q11.2 deletion syndrome (Amin et al. 2017) and the Fmr1 deficient model of fragile X syndrome (He et al. 2018). In the latter study, early postnatal correction of GABA depolarization (bumetanide-treated) led to sufficient normalization of the mature BC network (He et al. 2018). Therefore, the NKCC1 and KCC2 chloride transporters have been proposed as a potential therapeutic target of epilepsies by many studies in animal models and human patients (Moore et al. 2017).

Our study focuses in understanding the early developmental cellular and physiological mechanisms of mPFC circuits, before adolescence, and proposes that the neonatal mPFC compared to BC exhibits a delayed switch from depolarization to hyperpolarization function of GABAAR. Our results raise the possibility that the delayed maturation of mPFC compared to other cortical areas depends on a combination of a delayed switch from depolarization to hyperpolarization function of the GABAAR and delayed maturation of interneurons.

## Acknowledgements

Authors are grateful to Emmanuella Foinikianaki for her help with Matlab analysis and histology and to Giasemi Eptaminitaki for her help in *in situ* hybridization experiments. They also would like to thank all the members of Karagogeos and Sidiropoulou Labs and the animal facility of the IMBB for help with experiments.

This study was co-financed through the Operational Program “Education and Lifelong Learning” of the National Strategic Reference Framework – Research Funding Program (EDBM34) by a grant to DK (10040) and through the BIOIMAGING-GR, National Roadmap for Research Infrastructures from the European Union (European Social Fund-ESF) and Greek National Funds. KK has been a recipient of the Manasaki fellowship and a Medical School fellowship of the University of Crete and a poster award at the 27th Hellenic Society for Neuroscience Meeting.

## Author Contributions

All experiments were conceived and designed by K.K., K.S., and D.K. All experiments performed by K.K., A.V., and O.C. Computational simulations were performed by K.S. Data were analyzed by K.K., A.V., O.C., M.D. K.S. and discussed with D.K. Manuscript was written by K.K., D.K. and K.S. All authors discussed and commented on the manuscript.

## Abbreviations

BC: Barrel Cortex
INs: Interneurons
KCC2: K-Cl co-transporter 2
NKCC1: Na^+^-K^+^-Cl^−^ cotransporter 1
mPFC: medial Prefrontal Cortex
P10: Postnatal day 10
P20: Postnatal day 20
PNs: Pyramidal neurons
sEPSC: spontaneous excitatory postsynaptic current
sIPSC: spontaneous inhibitory postsynaptic current
YFP: yellow fluorescent protein
PV: parvalbumin
SST: somatostatin

