## Supplemental figures for "The developmental changes in intrinsic and synaptic properties of prefrontal neurons enhance local network activity from the second to the third postnatal week in mice"

### Supplementary information

#### Supplementary Figure 1: diazepam effect in BC at p10 and p20

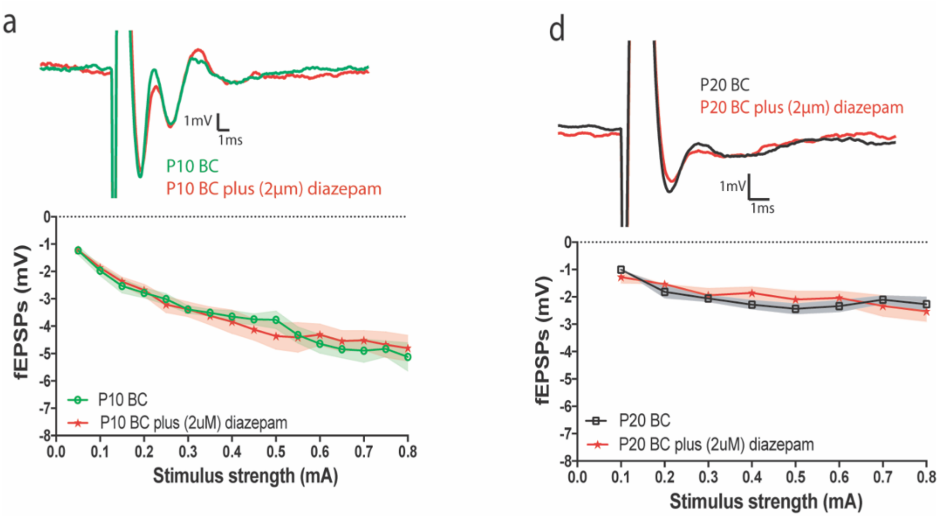

**(a)** Graph (bottom) and representative traces (top) showing that diazepam bath application does not have any effect on the fEPSP amplitude in BC at P10. Two-way repeated measures ANOVA analyses of evoked fEPSPs revealed a significant effect of stimulus strength (F (15, 140) = 24.05, p<0.0001) but not experimental conditions ($F_{(1, 135)}$ = 0.03, p=0.86), (n=6-7 brain slices from 3-4 mice).

**(b)** Graph (bottom) and representative traces (top) showing that bath application of diazepam does not have any effect in the fEPSP amplitude in BC at P20. Two-way repeated measures ANOVA analyses of evoked fEPSPs revealed a significant effect of stimulus strength ($F_{(7, 96)}$ = 5.51, p<0.0001) but not experimental conditions ($F_{(1, 96)}$ = 0.50, p=0.47), (n=6-7 brain slices from 3-4 mice).

**Supplementary Figure 2: No cell death contribution, at P10 and P20, in the establishment of mPFC IN population number.**

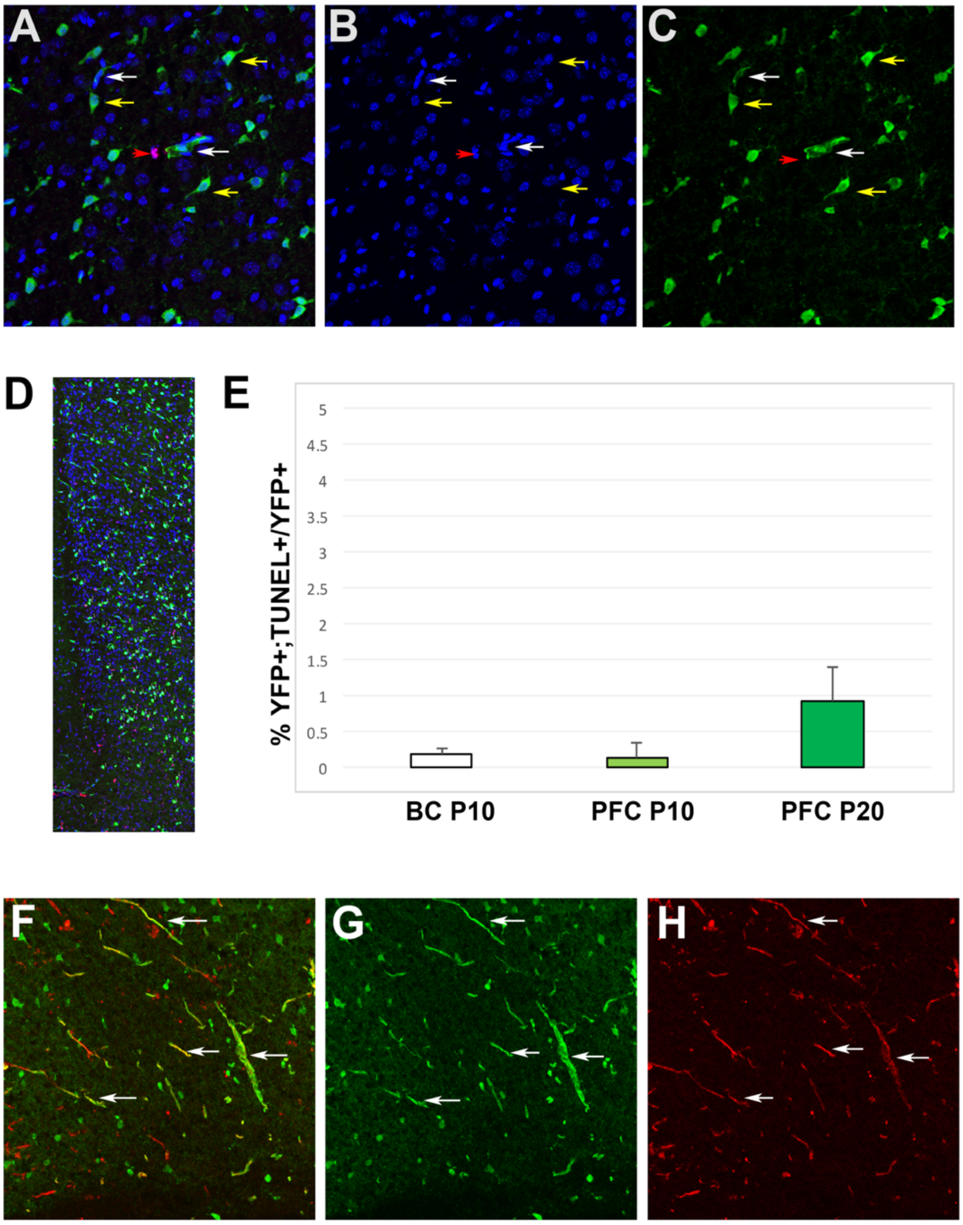

**(a-c)** Representative photos of the immunohistochemistry performed. Green=$YFP^+$; $Lhx6^+$ INs (yellow arrows) and $YFP^+$; $Lhx6^+$ epithelial blood vessel cells (white arrows). Blue=DAPI. RED=TUNEL, dead cell (red arrow)

In our counting we focused on the $YFP^+$; $Lhx6^+$ IN population

**(d)** Representative photo for our counting

(d) Graph of our counting results

**(f-h)** Immunohistochemistry for YFP (green, G), and CD31 (red; marker of endothelial blood vessel cells, H). Arrows show the blood vessels.

**Supplementary Figure 3: PV⁺ cells were not found in mPFC p10 but were identified in BC, at P10.**

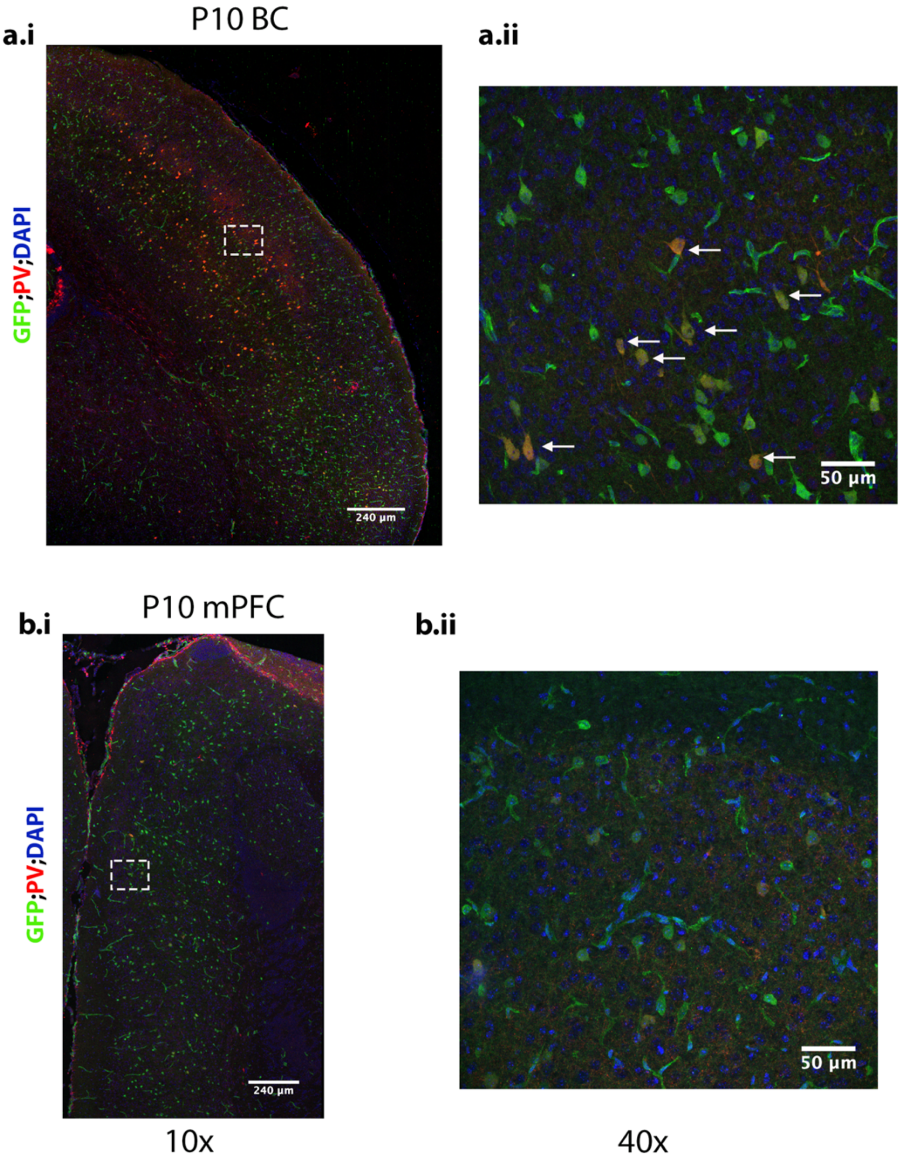

Representative triple immunostaining for GFP; PV (PV: parvalbumin) and DAPI in mPFC and BC at P10 is shown. $PV^+$ cells were not found in mPFC (b) but were identified in BC, at P10. Arrows in a.ii show the presence of cells stained for DAPI, GFP and PV. No such cells could be identified in b.ii.

**Supplementary Figure 4: Differential development of total cell density.**

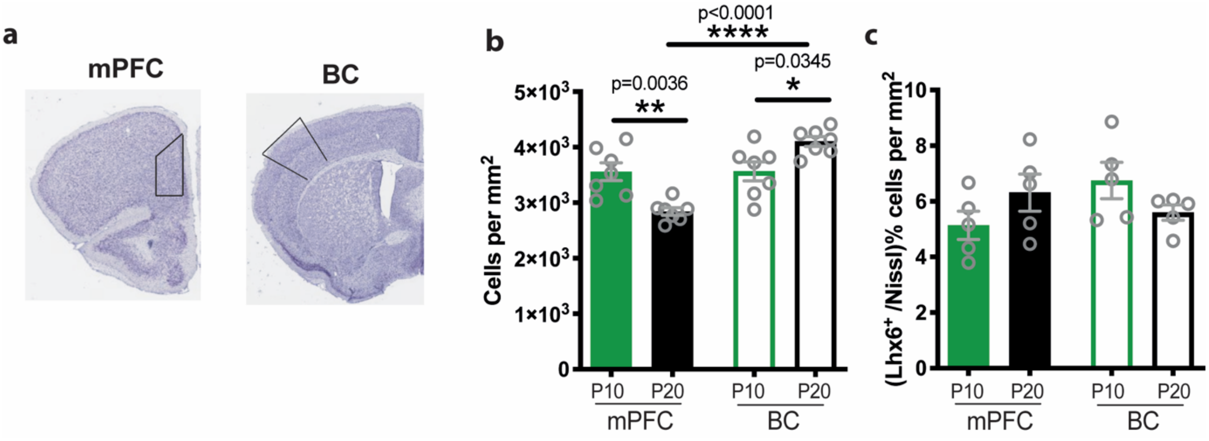

**(a)** Coronal sections with Nissl staining at P20 of mPFC and BC. The boxes (outlines) indicate the area that was measured in mPFC and BC.

**(b)** Bar graph showing the number of cells per $mm^2$ from Nissl staining (total cell density) during development at P10 and P20, in the mPFC and BC. Two-way ANOVA analyses of the cell density revealed a significant effect of brain area ($F_{(1,24}$=23.79, p<0.0001) but not of age ($F_{(1,24)}$=0.46, p=0.50). Post-hoc analysis showed that the number of cells per $mm^2$ is significantly lower in mPFC compared to BC, at P20 (Tukey's test, p<0.0001). In addition, the number of cells per $mm^2$ are significantly decreased at P20 compared to P10 in mPFC (Tukey's test, p=0.0036), but increased at P20 compared to P10 in BC (Tukey's test, p=0.0.0345), (n=7 mice/group).

**(c)** Bar graph showing the total cell density of $Lhx6^+$ neurons over to total cell density (Nissl positive cells) of mPFC and BC at P10 and P20. Two-way ANOVA analyses showed no significant effect of age ($F_{(1,\ 16)}$ = 0.000334, p=0.99) or brain area ($F_{(1,\ 16)}$ = 0.65, p=0.43) was found (n=5 mice/age group).

**Supplementary Figure 5: Changes in the passive properties of mPFC and BC pyramidal neurons with age.**

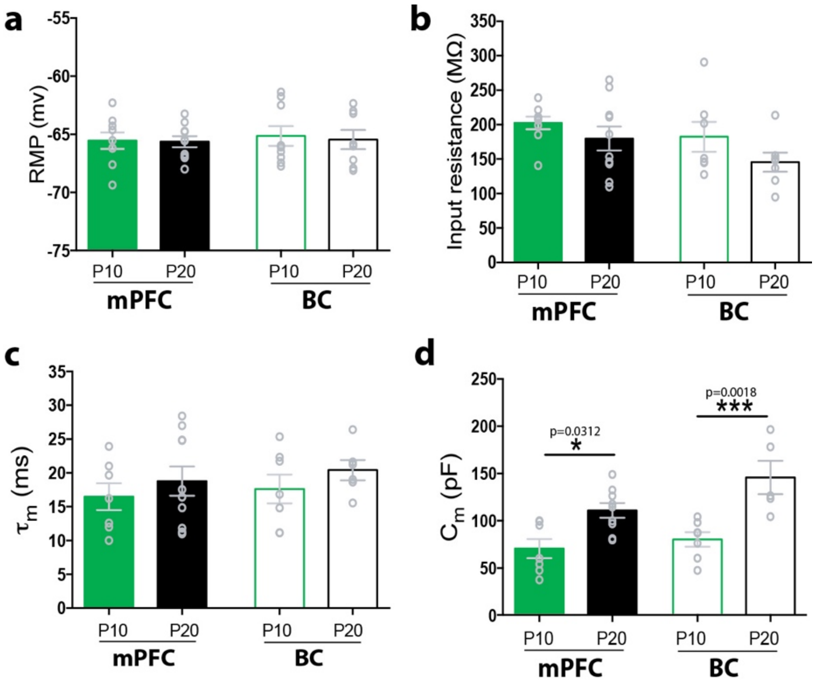

**(a)** Bar graph showing the resting membrane potential (RMP) of layer II/III pyramidal neurons at P10 and P20 in mPFC and BC. Two-way ANOVA analyses showed no significant effect of age ($F_{(1,32)}$ = 0.074, p=0.78) or brain area ($F_{(1,32)}$ = 0.16, p=0.68) was found (n=6-9 cells from 6-10 mice/age group).

**(b)** Bar graph showing the input resistance of layer II/III pyramidal neurons at P10 and P20 in mPFC and BC. Two-way ANOVA analyses showed no significant effect of age ($F_{(1,26)}$ = 3.65, p=0.067) or brain area ($F_{(1,26)}$= 2.16, p=0.15) was found (n=8-9 cells from 6-10 mice/age group).

**(c)** Bar graph showing the membrane time constant (τm) of layer II/III pyramidal neurons from P10 to P20 in mPFC and BC. Two-way ANOVA analyses showed no significant effect of age ($F_{(1, 26)}$ = 1.44, p=0.23) or brain area ($F_{(1, 26)}$= 0.42, p=0.51) was found (n=8-9 cells from 6-10 mice/age group).

**(d)** Bar graph showing the membrane capacitance (Cm) of layer II/III pyramidal neurons from P10 to P20 in mPFC and BC. Two-way ANOVA analyses showed a significant effect of age ($F_{(1,24)}$ = 26.03, p<0.0001) and brain area ($F_{(1,24)}$= 0.99, p=0.042) was found. Post-hoc analysis showed that Cm significantly increased at P20 compared to P10 in mPFC and BC, respectively (Tukey's test, p=0.031 and p= 0.0018, respectively), (n=6-9 cells from 6-10 mice/age group).

**Supplementary Figure 6: Input-output curves of mPFC and BC pyramidal neurons at P10 and P20.**

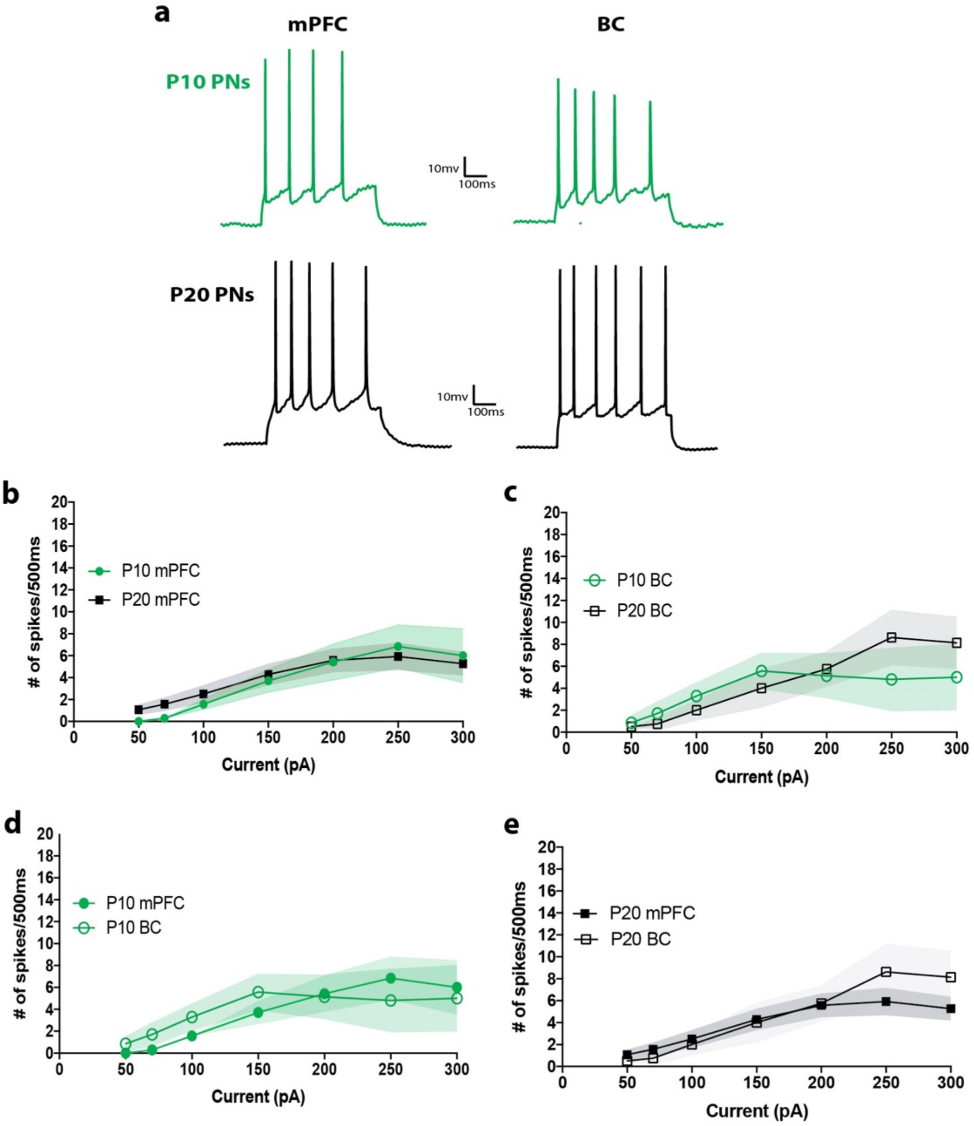

**(a)** Representative traces from mPFC (left) and BC (right) at P10 (green) and P20 (black).

**(b)** Graph showing no difference in the number of spikes per 500ms of increasing current stimulation at P10 and P20 in mPFC. Two-way ANOVA analyses revealed a significant effect of current stimulation ($F_{(6, 126)}$ = 8.89, p<0.0001) but not ages ($F_{(1, 126)}$ = 0.3, p=0.58) was found, (n=9-14 cells from 6-10 mice/age group).

**(c)** Graph showing no difference in the number of spikes per 500ms of increasing current stimulation at P10 and P20 in BC. Two-way ANOVA analyses revealed a significant effect of current stimulation ($F_{(6, 86)}$ = 4.11, p=0.0011) but not between ages ($F_{(1, 86)}$ = 0.28, p=0.60) was found, (n=8-9 cells from 6-7 mice/age group).

**(d)** Graph showing no difference in the number of spikes per 500ms of increasing current stimulation between mPFC and BC, at P10. Two-way ANOVA analyses revealed a significant effect of current stimulation ($F_{(6, 78)}$ = 4.12, p=0.0012) but not between brain areas ($F_{(1, 78)}$ = 0.18, p=0.66) was found, (n=8-9 cells from 6-7 mice/age group).

**Supplementary Table 1. Intrinsic electrophysiological properties of pyramidal neurons and interneurons in mPFC and BC at P10 and P20.**

| Interneurons | | | | |
|---|---|---|---|---|
| | P10 mPFC | P20 mPFC | P10 BC | P20 BC |
| Resting membrane potential, RMP (mv) | -64.81 ± 2.11, n=6 | -62.22 ± 0.94, n=8 | -63.29 ± 1.54, n=9 | -60.92 ± 3.38, n=6 |
| Input Resistance (MΩ) | **391.7 ± 73.62, n=8** | 129.2 ± 23.91, n=8 | 159.8 ± 17.89, n=9 | 144.2 ± 10, n=6 |
| Membrane time constant, τm (ms) | **36.94 ± 8.18, n=8** | 20.72 ± 8.37, n=6 | 10.29 ± 2.45, n=8 | 4.597 ± 0.73, n=6 |
| Membrane Capacitance, Cm (pF) | **80.9 ± 10.09, n=6** | 272.6 ± 109.8, n=6 | 49.2 ± 3.67, n=7 | 28.02 ± 3.6, n=6 |
| AP amplitude (mv) | **78.92 ± 4.15, n=6** | 76.41 ± 8, n=8 | 57.56 ± 3.84, n=8 | 79.60 ± 3.85, n=4 |
| Rate of rise of AP: dV/dt (mv/ms) | 90.49 ± 3.7, n=6 | 166.7 ± 21.48, n=6 | **66.41 ± 4.24, n=8** | 201.90 ± 22.71, n=4 |
| Duration of AP: Half-width (ms) | 2.24 ± 0.17, n=6 | **1.31 ± 0.15, n=8** | 2.49 ± 0.23, n=7 | 0.92 ± 0.19, n=4 |
| Rheobase (pA) | 170 ± 20, n=5 | 150 ± 15.43, n=7 | 162.5 ± 15.67, n=8 | 132.5± 31.19, n=4 |
| Threshold (mv) | -61.52 ± 1.5, n=5 | -56.58 ± 2.94, n=8 | -54.79 ± 3, n=9 | -50.07 ± 1.9, n=4 |
| AHP amplitude (mv) | -4.65 ± 0.95, n=4 | -5.96 ± 0.72, n=6 | **-13.99 ± 1.69, n=7** | 20.51 ± 1.97, n=4 |
| AHP time (ms) | 14.64 ± 1.29, n=5 | 13.26 ± 3.07, n=7 | 16.57 ± 2.47, n=7 | 18.63 ± 7.05, n=4 |

± represents the standard error of mean and n represents the number of cells. With **bold** presents the significant differences after post-hoc analysis (more details in figure legends).
